# Insecticide resistance of *Culex pipiens* f. *pipiens* is geographically widespread in Germany and displays an unmatched genotypic and phenotypic resistance state

**DOI:** 10.64898/2026.09.23.753539

**Authors:** Emma Vandenberghe, Stien Vereecken, Friederike Reuss, Isabell Brusius, Corneel Janssens, Axel Magdeburg, David A. Groneberg, Ruth Müller, Isabelle Marie Kramer

## Abstract

In Germany, insecticide use is recommended during mosquito-borne disease outbreaks, yet nationwide data on insecticide resistance in key vector species are scarce. This is particularly relevant for *Culex pipiens* s.l., a vector of West Nile virus. Here, we provide the first broad-scale assessment of molecular insecticide resistance (kdr and ace-1 mutations) in *Cx. pipiens* f. *pipiens* across Germany. We associated insecticide resistances with environmental conditions, potential links with mitochondrial variation of the cytochrome c oxidase I (COI) gene as well as phenotypic resistance (CDC bottle assay with permethrin) of the Frankfurt population with elevated kdr frequencies.

Kdr resistance alleles were detected in German *Cx. pipiens* f. *pipiens* reaching frequencies of up to 50% in the metropolitan areas of Berlin and Frankfurt am Main. Annual precipitation was negatively associated with kdr allele frequency. Interesstingly, ace-1 resistance mutations were only present in the metropolitan area of Frankfurt am Main and in Muggensturm (Rhine valley, southwestern Germany). COI haplotype analyses revealed low genetic variability within German *Cx. pipiens* f. *pipiens* populations with no association to the insecticide resistance genotype. Phenotypic assays of the Frankfurt population showed early signs of permethrin resistance, but unmatched genotypic and phenotypic resistance state.

This study provides the first broad-scale assessment of molecular insecticide resistance in *Cx. pipiens* f. *pipiens* across Germany, establishing a baseline for resistance surveillance and vector control. Further research is needed to identify factors shaping phenotypic resistance, as molecular markers alone may not reliably predict resistance. Since sampling was restricted to urban green spaces, broader sampling across environmental gradients is needed to identify additional environmental effectors.

## Background

In Europe, climate warming is driving the expansion of West Nile virus (WNV), transmitted by native *Culex* mosquitoes (Fay et al., 2025), posing a major mosquito-related public health concern alongside the spread of the Asian tiger mosquito (Fay et al., 2025). In 2025, nine countries in Europe reported 652 acquired human cases of WNV infections (ECDC, 2025). In 2018, Europe experienced its largest recorded WNV outbreak so far, with more than 2,000 human cases (Frank et al., 2022). During this outbreak, the virus was detected for the first time in birds and horses in Germany (Frank et al., 2022). Since then, the virus has caused recurrent seasonal outbreaks in Germany (Offergeld et al., 2025). In addition, nationwide monitoring surveys in Germany detected several mosquito-borne viruses (MBD), such as Sindbis virus, Usutu virus and Batai virus in field collected *Cx. pipiens* f. *pipiens*, *Cx. Pipiens* f*. molestus* and *Cx. torrentium*, confirming their role as important vectors in the country (Erazo et al., 2024).

Vector control is essential to reduce arboviral transmission risk, particularly in the absence of specific treatments or licensed vaccines against MBDs such as WNV (Wang et al., 2022). The control of mosquitoes is primarily based on environmental management, larviciding (e.g. *Bacillus thuringiensis israelensis*; Bti) and insecticide treatment of adults (Paaijmans et al., 2019). However, insecticide resistance is rising in Europe driven by exposure to previous vector control pesticides used in agriculture or at households (Arich et al., 2024). Phenotypic resistance to pyrethroids, dichlorodiphenyltrichloroethane (DDT) and in some cases bendiocarb have already been reported in Greece, Spain, Italy, and Belgium(Kioulos et al., 2014; Paaijmans et al., 2019; Pichler et al., 2022; Vereecken et al., 2022). In addition, frequencies of resistance alleles (e.g. voltage gated sodium channel domain II L1014F mutation (knockdown-resistance; kdr) and acetylcholinesterase 1 G119S (ace-1)) have increased, reaching for instance 88.3% in Greece (Csiba et al., 2025; Fotakis et al., 2017; Pichler et al., 2022; Wang et al., 2022). In Germany, insecticides are recommended in case of an MBD outbreak such as WNV (FLI, 2022), yet the resistance status of mosquitoes as well as environmental drivers of resistance, particularly in *Cx. pipiens*, remain largely unknown (Portwood et al., 2025), raising concerns about failure to effectively control mosquito populations during outbreaks.

To generate baseline insecticide resistance data across Germany and support preparedness for potential insecticide use during outbreaks, we conducted the first comprehensive assessment of molecular insecticide resistance in *Cx. pipiens f. pipiens* across Germany, examined potential environmental drivers of resistance-associated variants, and evaluated genetic COI variability to explore possible associations between COI haplotype and resistance genotype. In addition, we assessed genotype– phenotype relationships for insecticide resistance for one population by testing its phenotypic permethrin resistance and comparing these results with the populations’ molecular kdr resistance status. The resulting data are intended to fill critical knowledge gaps and to support future predictive modelling and vector management strategies. In detail, this study aimed to (i) map molecular insecticide resistance across German *Cx. pipiens* f. *pipiens* populations using kdr and ace-1 markers and analyse how resistance-allele frequencies relate to environmental variables, (ii) investigate COI genetic variability in *Cx. pipiens* f. *pipiens* populations across Germany to identify potential COI haplotype associations with molecular resistance genotypes, and (iii) verify phenotypic and molecular insecticide resistance in a population to establish genotype–phenotype resistance patterns (Figure 1). Likewise, it was hypothesised that (i) kdr and ace-1 resistance allele frequencies are heterogeneously distributed across German *Cx. pipiens* f. *pipiens* populations and are positively associated with anthropogenic factors (Arich et al., 2024; Vereecken et al., 2022); (ii) *Culex pipiens* f. *pipiens* populations across Germany exhibit low genetic variability at the COI locus and COI haplotype cannot be linked to a specific resistance genotype (Werblow et al., 2014); (iii) populations with elevated molecular resistance markers exhibit corresponding phenotypic resistance supporting a clear genotype–phenotype relationship (Scott et al., 2015).

**Figure 1:**
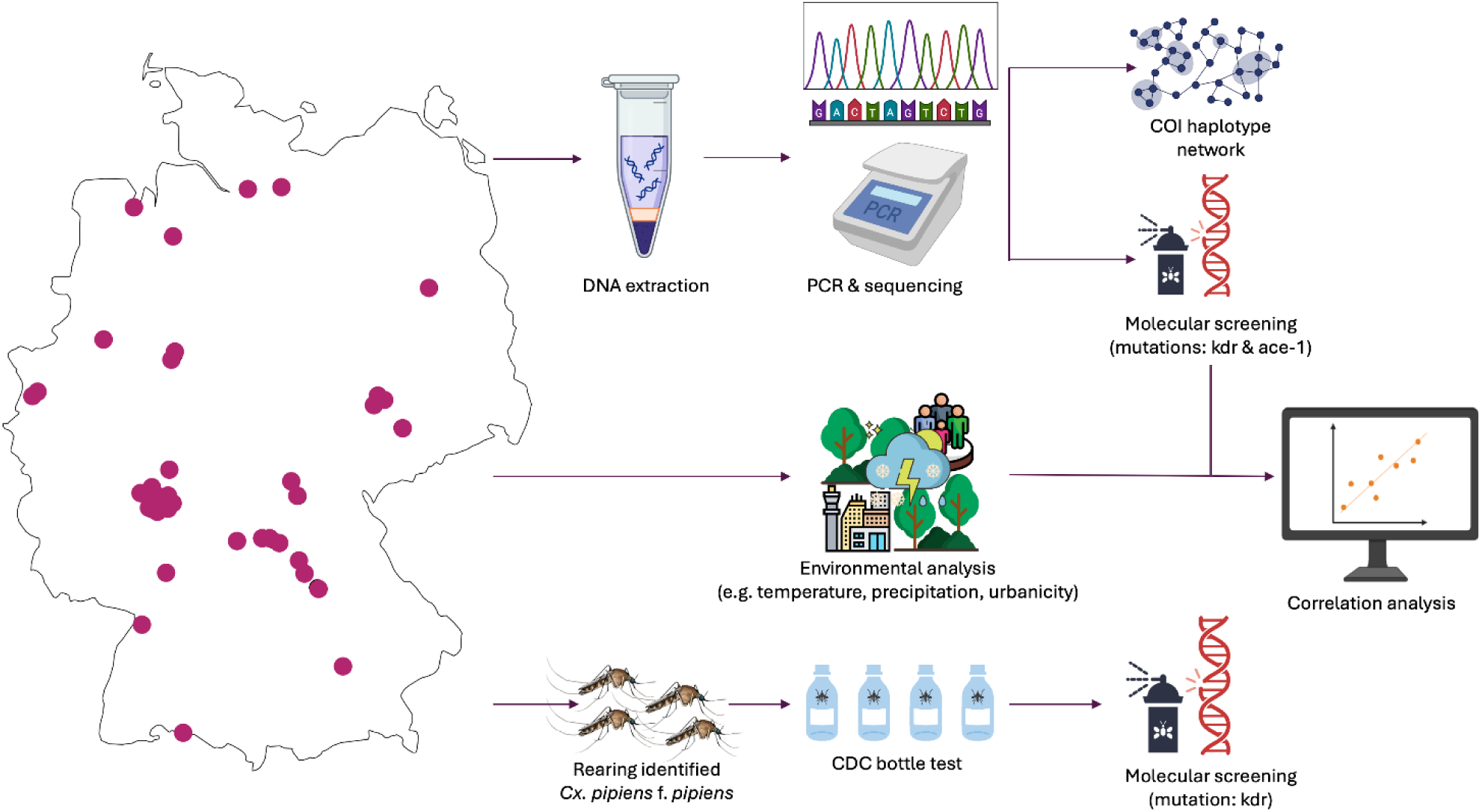
Study design for identification and analysis of molecular and phenotypic insecticide resistance in *Culex pipiens* f*. pipiens*. A subset of *Cx. pipiens* f. *pipiens* individuals were used for COI genetic analyses to capture potential mitochondrial differences among the different populations across Germany. A subset of the same species was also subjected to kdr and ace-1 variant detection to determine the molecular insecticide resistance status. The results of the molecular insecticide resistance screening were associated to environmental variables of every sampling site. Lastly, a population of *Cx. pipiens* f. *pipiens* from Frankfurt am Main was used for assessing phenotypic resistance to permethrin with the CDC bottle test and for molecular insecticide resistance screening of the kdr mutation frequency. Figure created with BioRender.com.

## Materials and methods

### Study area and environmental description of the sampling sites

In 2023, 41 locations were sampled for larvae and pupae across Germany (Supplementary Figure 1, Supplementary Table 1). Sampling was conducted by both scientists and citizen participants. Sampling locations consisted of urban green spaces located near human settlements. The climate at each sampling site was analysed focusing on air temperature (°C), relative humidity (%) and rainfall (mm). Hourly climatic data was extracted from the Historical Weather API for the sampling year, respectively from January 1 to December 31, 2023 (Zippenfenig, 2023). In addition to the annual climatic variables, the high-resolution bioclimatic variables (BIO1-BIO19) were derived to further characterize the mosquito sampling sites, representing a global, kilometer-scale climate dataset generated with WorldClim Version 2 (1970-2000) (Fick & Hijmans, 2017). Environmental variables linked to urbanisation such as built-up surface/10,000 m^2^, population density (people/10,000 m^2^) and settlement (no unit, values represent urbanisation classes) were extracted from the Global Human Settlement Layer of the year 2020 using the Mollweide coordination system (Ehrlich et al., 2018; GHS Manual, 2023). The enhanced vegetation index (EVI) was extracted from the MODIS Collection 6 (MOD13A3) (Didan & Barreto Munoz, 2019). The generated and downscaled MOD13A3_061 1_km_monthly_EVI values were used to calculate a mean value per sampling site. The means of the environmental parameters were graphically visualized per sampling site using R (version 2025.05.0) and RStudio (version 2025.05.0). In the subsequent analyses, the R packages *terra* (v4.3.3), *dplyr* (v1.1.3), *readr* (v2.1.4), *janitor* (v4.3.3), and *tidyr* (v1.3.0) were used (Supplementary Figures 2-4).

### Sample collection and preparation for bioassays

At each site at least three randomly selected breeding habitats were searched for 20 larvae or pupae each with the aim to sample *Culex*. Mainly graveyards and gardens were sampled, but also botanical gardens and green village squares were included. Breeding habitats were mostly grave vases and pouring water pools, but the following 18 other habitat types were sampled: Watering can, mortar bucket, rain barrel, bucket, artificial pond, flowerpot, grave vase with lid, clogged drain, milk churn, bird bath, bathtub, oviposition trap, trough, tractor tires, spring, stone ornament, drain, and flowerpot holder. The larvae and pupae were stored in 70% ethanol until further processing.

In addition, egg rafts were sampled in September 2023 and summer 2024, and included in the COI haplotype analysis. *Culex* egg rafts were collected using eight oviposition traps which consist of black plastic containers of multiple forms and sizes, filled with rainwater and dried horsetail. Egg rafts were sampled at Berlin Mitte (graveyard), Berlin-Marienfelde (green area in front of a building), Rietschen (private garden), Frankfurt-Westend (Botanical Gardens Frankfurt), Waldems-Reinborn (graveyard, gardens) and Schotten-Breungeshain (graveyard, gardens).

In 2025, egg raft samples for genotypic and phenotypic resistance tests were collected at the Botanical Gardens in Frankfurt am Main (Germany). The egg rafts were transferred to 1.5 ml Eppendorf tubes and kept at fridge temperature (4°C to 8°C), until the hatching. Eighty-five egg rafts were reared at 27°C, 80% relative humidity and 16:8 hours light dark rhythm in climatic cupboards (CPS-P530 Climatic Cabinet, RUMED, Germany). Each egg rafts was separately hatched in 100 ml plastic cups filled with softened water and crushed TetraMin fish food (Tetra, Melle, Germany). After 48 hours, five larvae per egg raft were pooled and DNA was extracted. Two hundred to 400 larvae of *Cx. pipiens* f. *pipiens* were placed together in a plastic rearing tray with 800 ml of softened water and TetraMin *ad libitum*. All female pupae were collected in a 100 ml cup and placed into a 17 cm x 17 cm rearing cage for emergence, provided with a 10 % glucose solution. Molecular identification was performed of each egg raft as described below and solely *Cx. pipiens* f. *pipiens* was used for further analyses (bioassay for phenotypic permethrin resistance and molecular PCR analysis for genotypic kdr mutation).

### Molecular species identification

The DNA extraction was performed using the Phire Animal Tissue Direct PCR kit (Thermo Fisher Scientific, Waltham, USA), according to the manufacturer’s protocol. While immature stages were used individually in the subsequent analyses, five L1 larvae per egg raft were pooled for DNA extraction for species barcoding. The CQ11 marker was used to distinguish between *Cx. pipiens* f. *pipiens* and *Cx. pipiens* f. *molesutu*s based on fragment sizes and *molestus*/*pipiens* interform hybrids by double bands (Danabalan et al., 2012). For the individuals identified as *Cx. pipiens* s.l. the acetylcholinesterase-2 (ace-2) marker was used to rule out sibling species *Cx. torrentium*. The CQ11 microsatellite locus was amplified using the primers CQ11F2 (5’-GATCCTAGCAAGCGAGAAC-3’), MOLCQ11R (5’-CCCTCCAGTAAGGTATCAAC-3’) and PIPCQ11R (5’-CATGTTGAGCTTCGGTGAA-3’) (Bahnck & Fonseca, 2006). The 20 μL PCR reaction mixture contained 1 μL DNA template, primers at final concentrations of 25 pmol/μL, AccuGENE H_2_O and GOTAQ G2 hotstart polymerase (Promega, Madison, USA). The PCR consisted of two sequential cycling procedures, of which the conditions of the first cycling were the following: Initial activation step at 95°C for 2 minutes; 10 cycles of denaturation at 95°C for 30 seconds, annealing at 60°C for 30 seconds and elongation at 72°C for 30 seconds. The second cycling followed 25 cycles of denaturation at 95°C for 30 seconds, annealing at 54°C for 30 seconds and elongation at 72°C for 30 seconds, a final elongation at 72°C for 5 minutes and a cool down at 16°C. The ace-2 gene was amplified using the forward primer B1246s (5’-TGGAGCTCCTCTTCACGG-3’) and the reverse primers ACEpip (5’-GGAAACAACGACGTATGTACT-3’) and ACEtorr (‘5-TGCCTGTGCTACCAGTGATGTT-3’) (Smith & Fonseca, 2004), to distinguish between *Cx. pipiens* s.l. and *Cx. torrentium* (Farajollahi et al., 2011). The 20 μL PCR reaction mixture contained 1 μL DNA template, primers at 10 pmol/μL final concentration, AccuGENE H_2_O and GOTAQ G2 hotstart polymerase (Promega, Madison, USA). The cycle started with an initial activation at 95°C for 2 minutes. Followed by 40 cycles of denaturation at 95°C for 30 seconds, annealing at 50°C for 45 seconds and elongation at 72°C for 60 seconds. Then a final elongation step at 72°C for 10 minutes and lastly a cool down at 15°C.

All PCR products were checked on a 2% agarose gel, at 100V for 35 (COI), 40 (ace-2) or 60 (CQ11) minutes. Before separation, GelRed nucleic acid gel stain (Biotium, Fremont, USA) was added to the agarose mixture for visualisation with the GelDoc XR+ (BioRad, Hercules, USA). All field specimens that could not be identified by ace-2 and CQ11 fragment size analyses were subjected to forward cytochrome c oxidase subunit I (COI) sequencing to determine species identity. The COI gene fragment was amplified using the barcoding primers LCO 1490 (5’-GGTCAACAAATCATAAAGATATTGG-3’) and HCO 2198 (5’-TAAACTTCAGGGTGACCAAAAAATCA-3’) (Folmer et al., 1994). The 20 μL PCR reaction mixture contained 1 μL DNA template, both primers at 25 pmol/μL concentrations, AccuGENE H_2_O and 60TAQ G2 hotstart polymerase (Promega, Madison, USA). The cycle started with an initial activation at 95°C for 2 minutes. Followed by 40 cycles of denaturation at 95°C for 30 seconds, annealing at 50°C for 45 seconds and elongation at 72°C for 60 seconds. Then a final elongation step at 72°C for 10 minutes and lastly a cool down at 15°C for 10 minutes.

All COI sequences were analyzed using Geneious Prime 2025.0.3 (https://www.geneious.com) and manually checked to ensure sequence quality. Thereafter, the individuals were identified, with a reliability of >98%, using nBLAST® from the National Library of Medicine (https://blast.ncbi.nlm.nih.gov/Blast.cgi).

A subset of *Cx. pipiens* f. *pipiens* individuals, randomly chosen while ensuring that as many sampling sites as possible were represented, were subjected to forward and reverse COI sequencing, to identify haplotype diversity of *Cx. pipiens* f*. pipiens* (Figure 5, Supplementary Table 3). Haplotype analysis was conducted on 84 consensus sequences. The consensus sequences were aligned with the Clustal Omega algorithm in Geneious Prime 2025.0.3 (Sievers et al., 2011). Haplotypes were identified with DnaSNP Version 6 (Rozas et al., 2017). Thereafter, two haplotype networks were created in PopART (Leigh & Bryant, 2015) using the TCS method (Templeton et al., 1992). In addition, the haplotype diversity and nucleotide diversity were calculated in R using the packages *ape* and *pegas* (Grünwald & Hoheisel, 2006; Paradis & Schliep, 2019). To identify how the genetic variation was partitioned among the populations, an AMOVA statistical test was performed using the R packages *ape, pegas, adegenet* and *poppr* (Grünwald & Hoheisel, 2006; Jombart, 2008)

### Molecular insecticide resistance markers

A subset of *Cx. pipiens* f. *pipiens* were used to investigate the voltage gated sodium channel (VGSC) domain II for the presence of the L1014F mutation (knockdown-resistance; kdr). The acetylcholinesterase 1 (ace-1) gene was assessed for the G119S mutation, both mutations are linked to insecticide resistance in Culicidae (Martinez-Torres et al., 1999; Weill et al., 2004).

The kdr PCR requires two PCR reaction runs in parallel using four primers: Cgd1 (5ʹ-GTGGAACTTCACCGACTTC-3ʹ), Cgd2 (5ʹ-GCAAGGCTAAGAAAAGGTTAAG-3ʹ), Cgd3 (5ʹ-CCACC GTAGTGATAGGAAATTTA-3ʹ), and Cgd4 (5ʹ-CCACCGTAGTGAT AGGAAATTTT-3ʹ) (Martinez-Torres et al., 1999). The first PCR Mastermix consisted of 1 μL DNA template, AccuGENE H_2_O, GoTAQ G2 hotstart polymerase (Promega, Madison, USA) and the primers Cgd1, Cgd2 and Cgd3 (25 pmol/µl) needed to identify the susceptible allele. The second PCR Mastermix consisted of the primers Cgd 1, Cgd2 and Cgd4 (25 pmol/µl), needed to identify the resistant allele. In both PCR reactions the individuals should have an internal control band at 500 bp amplified with primers Cgd1 and Cgd2, which represented the gene wherein the mutation is present. When an additional band at 350 bp was present in the first PCR reaction with the primer Cgd3, it indicated the presence of the susceptible allele. When an additional band at 350 bp was present in the second PCR reaction with the primer Cgd4, it indicated the presence of the resistant allele. The results of both PCRs were combined and used to identify the genotype of every individual. The cycling conditions were as followed: An initial activation step at 95°C for 2 minutes; 40 cycles of denaturation at 94°C for 30 seconds, annealing at 48°C for 30 seconds and elongation at 72°C for 60 seconds; a final elongation step at 72°C for 10 minutes and lastly a cool down at 15°C. GelRed nucleic acid gel stain (Biotium, Fremont, USA) was added to the agarose mixture for visualization with the GelDoc XR+ (BioRad, Hercules, USA).

Ace-1 was amplified using the primers CxEx3dir (5ʹ-CGACTCGGACCCACTGGT-3ʹ) and CxEx3rev (5ʹ-GTTCTGATCAAACAGCCCCGC-3ʹ) (Wang et al., 2022). The 20 μL PCR reaction mixture contained 1 μL DNA template, primers in 25 pmol/µl concentration, AccuGENE H_2_O and GoTAQ G2 hotstart polymerase (Promega, Madison, USA). The cycling conditions were the following: An initial activation at 95°C for 2 minutes; 40 cycles of denaturation at 94°C for 30 seconds, annealing at 54°C for 30 seconds and elongation at 72°C for 60 seconds; a final elongation at 72°C for 10 minutes and lastly a cool down at 15°C. Afterwards, the amplicons were digested with the restriction enzyme AluI (Thermo Fisher Scientific, Waltham, USA). The mutation, a substitution from GGC to AGC, creates a recognition site for the restriction enzyme, resulting in two DNA fragments, which can be visualized on an 2% agarose gel. A 20 μL mastermix for the restriction was made with AccuGENE H_2_O, 10x Buffer Tango and 1 μL AluI enzyme. Ten μL amplicon of the first PCR was added to the 20 μL AluI Mastermix. Then, the samples were incubated at 37°C for 8 hours. When only one gene fragment is observed the mutation is not present, two gene fragments indicate resistance, and three gene fragments indicate that the mutation is present on only one allele (Weill et al., 2004). All PCR products were checked on a 2% agarose gel run at 100V for 40 minutes. GelRed nucleic acid gel stain (Biotium, Fremont, USA) was added to the agarose mixture for visualization with the GelDoc XR+ (BioRad, Hercules, USA).

### Associations between molecular insecticide resistance and environmental variables

Relationships between molecular insecticide resistance and environmental variables were analysed in a two-step framework. First, an exploratory screening was conducted to assess associations between kdr allele frequencies and individual environmental predictors. Continuous environmental variables were analysed using generalised additive models (GAMs) implemented in the R package *mgcv* (Wood, 2010) with smoothing terms fitted by restricted maximum likelihood (REML) (Morpurgo et al., 2024). This analysis was used to identify candidate predictors and to assess whether relationships were approximately linear or nonlinear. Functional form was classified based on the effective degrees of freedom (edf), with terms considered approximately linear at edf ≤ 1.2 and nonlinear at edf > 1.2. Categorical predictors, such as Settlement Model class (SMOD), were examined using linear models. After the GAM screening, Pearson correlation coefficients were calculated among environmental variables. Significant environmental predictors identified in the screening step were then assessed with respect to their interdependence, and only a reduced set of non-redundant candidate predictors was retained for subsequent modelling. Based on the exploratory screening and dependency testing, annual precipitation (BIO12), mean temperature of the warmest quarter (BIO10), and SMOD class were retained for targeted GAM analyses.

Secondly, targeted GAMs were fitted. Competing models were compared using Akaike’s Information Criterion (AIC) and corrected AIC for small sample size (AICc), and the best-supported model was selected based on the lowest AICc, ΔAICc, and model weights (Supplementary Tables 5-8).

Model diagnostics for the best-supported GAMs included basis-dimension checks using “*gam.check ()”*, assessment of concurvity, Shapiro-Wilk tests of residual normality, Breusch–Pagan tests for heteroscedasticity, and visual inspection of residual plots. Model performance was summarised using adjusted R² and percentage deviance explained. Data wrangling and visualisation were carried out using dplyr, tidyr, ggplot2, MuMIn, and lmtest (Barton, 2026; Wickham, 2009; Wickham et al., 2023, 2026; Zeileis & Hothorn, 2002). Ace-2 was not used in the analysis as it showed no significant correlations in Pearson correlation analyses.

### Phenotypic insecticide resistance of *Culex pipiens* f. *pipiens*

The CDC bottle bioassay was performed using permethrin-coated Wheaton Bottles with a total of 98 female mosquitoes in the test bottles and 29 female mosquitoes in the control bottle. (Centers for Disease Control and Prevention, 2024). To coat the inside of the exposure bottles, a permethrin stock solution containing the diagnostic dose of 43 μg/bottle was made by diluting 4.3 mg of permethrin in 100 ml of absolute ethanol (Centers for Disease Control and Prevention, 2024). The control bottle was coated with 1 ml of absolute ethanol, while the four exposure bottles were coated with 1 ml of the stock solution. The bottles were swirled, inverted and rotated to coat the bottom, the sides and the lid of the bottle, and dried with the lid off. On average 24 unfed females aged three to five days were added to the coated bottles. Mortality was recorded every five minutes until 15 minutes passed, then mortality was checked every 15 minutes until all mosquitoes were moribund. The treatment mortality was calculated as the proportion of dead mosquitoes relative to the total number of exposed individuals after the pre-determined threshold time of 30 minutes (Centers for Disease Control and Prevention, 2022). The population was considered resistant when the mortality was below 90%, as possible resistant when the mortality was 90%-96% and as sensitive when the mortality was 97%-100% (US Centers for Disease Control, 2022). Resistance at the diagnostic time was assessed visually using a mortality curve, which illustrates the relationship between exposure time and percentage mortality during the test. The curve was generated using the R packages ggplot2 and dplyr (Wickham, 2009; Wickham et al., 2023).

The correspondence between phenotypic and molecular kdr-based resistance was assessed by visually comparing survival rates per CDC test bottle with the proportion of kdr-positive individuals. Kdr mutations were identified via PCR as described above. Survival and kdr frequency were visualised using the R packages ggplot2, tidyr and dplyr (Wickham, 2009; Wickham et al., 2023, 2026).

## Results

### Sampling sites are shaped by environment

Mean air temperature ranged from 9°C to 13°C in the sampling year 2023. Maximum air temperatures where the highest in the southeast, while it was milder in the northwest. Regions in the northwest received more rainfall, while regions in the southeast were relatively dry (precipitation, rain, average humidity; Supplementary Figure 2). Especially sites in the Taunus Forest, Hesse, (798-825m) showed climatic extremes in temperature and precipitation (Supplementary Figure 2). The long-term bioclimatic variables showed similar patterns as the annual climatic variables across regions, but can exhibit different ranges, e.g., annual precipitation of 600-950 mm versus 800-1400 mm (Supplementary Figure 3). The bioclimatic variables gave additional insights into the climate, such as isothermality (BIO3), which varies across the sampling sites (Supplementary Figure 3). For temperature seasonality (BIO4), there was an increasing gradient from sampling locations in the northwest to the southeast (Supplement Figure 3). The EVI values ranged between 0.20-0.45. Lower EVI values were found in sampling locations near built-up areas, whereas higher values were found in sites secluded in natural surroundings (Supplementary Figure 4). The built-up surface variable varied from 0 m^2^, such as forests near Frankfurt (e.g. locations abbreviated NOR, WES) and Berlin (BE) to 3,893 m^2^ built-up area in Heidelberg (HD), where nearly 40% of the grid was covered with buildings. Population density was generally low, except for one region (Heidelberg; HD) where it was prominently higher (approx. 100 people/10 000m²). Major cities, such as Berlin (BE), Hamburg (HH), Frankfurt (NOR, WES), Bremen (HB) and Heidelberg (HD) showed the highest settlement values compared to more rural sites and smaller villages (Supplementary Figure 4).

Eighty-two % of sampled locations were cemeteries, followed by gardens (13%), botanical gardens (4%), and village squares (1%) (Figure 2A). Accordingly, grave vases were the most sampled breeding sites (45%), followed by pouring water pools (31%) and other container types (25%) (Figure 2A). Because most breeding sites were small, the volume of water in which larvae were found per site was limited (Figure 2A).

**Figure 2:**
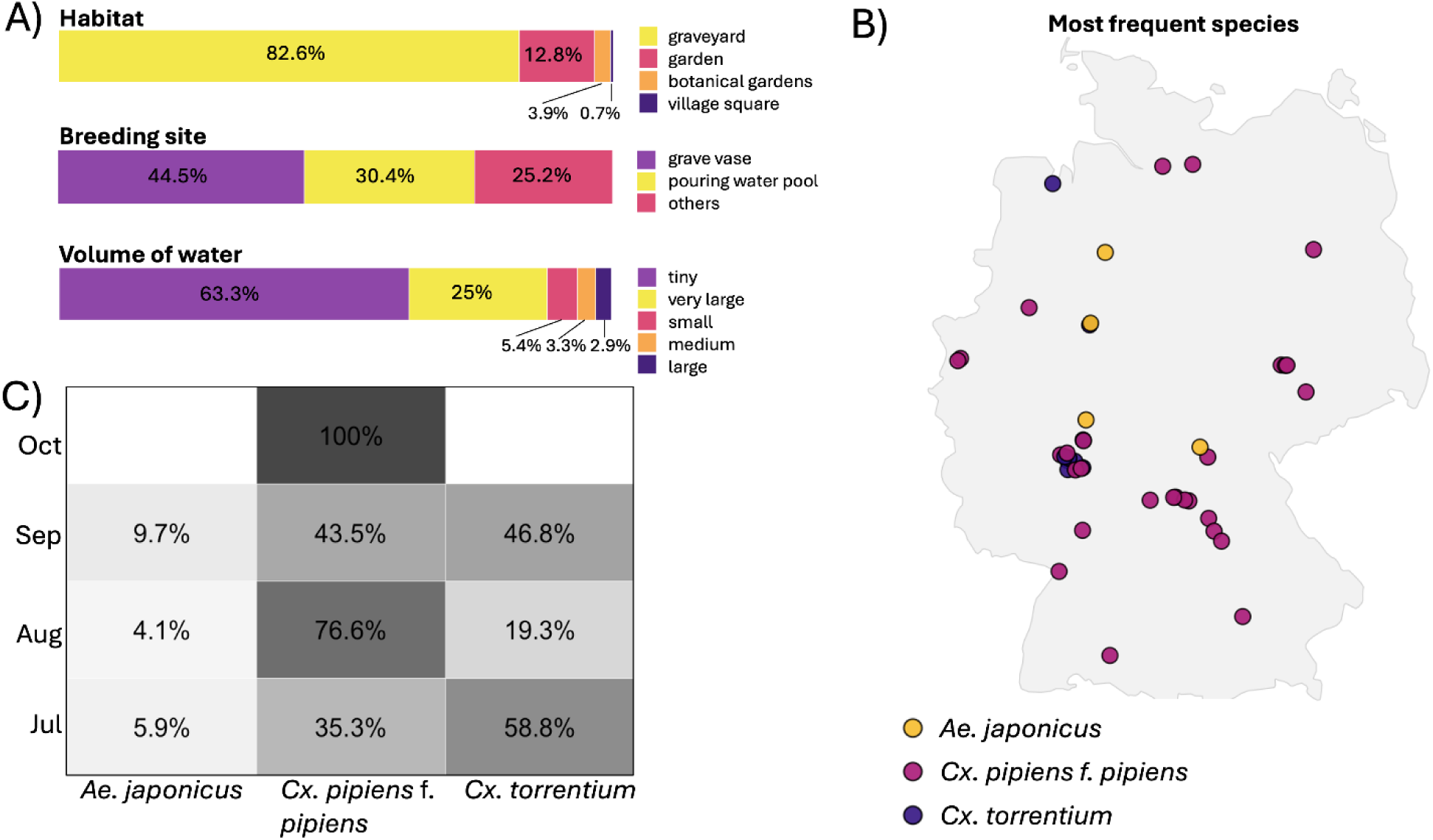
Habitat characteristics (A), species distribution across Germany (B), and seasonally most abundant species (C) in 2023. A) Description of breeding sites. Other breeding sites included watering cans, mortar buckets, rain barrels, buckets, artificial ponds, flowerpots, grave vases with lid, clogged drains, milk churns, bird baths, bathtubs, oviposition traps, troughs, tractor tires, springs, stone ornaments, drain and flowerpot holders. Categories of the water volume: tiny: <10L, small: 10-50L, medium: 50-100L, large: 500-1000L, very large: >1,000L. B) The most abundant mosquito species per sampling site. C) Seasonal distribution of the three most abundant species per sampling month.

### *Culex pipiens* f. *pipiens* is the most abundant mosquito species in urban green spaces

In total, 1,551 samples collected throughout 41 sampling locations across Germany were subjected to species identification. No species could be assigned in 344 samples, because of failed DNA extraction and/or PCR reaction (Supplementary Tables 1-3). Eight samples were identified as non-mosquito species, such as *Dasyhelea* spp., *Nannochloropsis limnetica*, *Lagenidium caudatum, Saprolegnia ferax* and *Cypridopsis vidua*. Thus, 1,199 samples were successfully identified: *Cx. pipiens* f. *pipiens* (59.5%) > *Cx. torrentium* (28.5%) > *Ae. japonicus* (5.5%) > *Cx. pipiens*/*molestus* hybrid (3.5%) > other mosquito species (1%). The latter including *Ae. geniculatus, Anopheles maculipennis, An. plumbeus* and *Cx. impudicus.* Ninety-five percent of all samples belonged to the three most abundant species, respectively *Cx. pipiens* f. *pipiens*, *Cx. torrentium* and *Ae. japonicus.* The highest number of five mosquito species per site was detected in Pohlheim (GI), Bad Lippspringe (PB_2) and Geiselwind (KT_2), respectively. In most of the sampling sites, two or three mosquito species were present. *Culex pipiens* f. *molestus* was detected in the 13 sampling sites in south-eastern and central Germany, namely, Erlangen (ER), Freising (FS), Pohlheim (GI), Glashütten (GLA), Heidelberg (HD), Hofheim-Lorsbach (HOF), Leipzig (LC), Bad Camberg (LM_1), Wechselburg (MW), Bad Lippspringe (PB_2), Frankfurt-Sossenheim (SOS), Weilrod-Riedelbach (WEI), and Frankfurt-Westend (WES) (Figure 2B, Supplementary Figure 1). Looking at co-occurrences, *Cx. pipiens* f. *pipiens* appeared together with *Cx. torrentium* and *Cx. pipiens/molestus* hybrids, while *Culex torrentium* co-occurred with *Cx. pipiens* f. *pipiens* and *Cx. pipiens* f. *molestus* (Supplementary Figure 5). The seasonal occurrence of the three most abundant mosquito species, *Cx. pipiens* f. *pipiens*, *Cx. torrentium* and *Ae. japonicus,* showed that *Culex torrentium* was the most abundant species in July and *Cx. pipiens* f. *pipiens* was the most abundant species in August (Figure 2C). In September, both *Culex* species were equally abundant, while in October only *Cx. pipiens* f. *pipiens* was detected (Figure 2C). *Culex pipiens* f. *molestus* was detected from July to October at low abundances with the highest frequency of occurrence observed in August (Figure 2C).

### Molecular insecticide resistance is widely distributed

The kdr mutation was detected at 22 of the 38 sampling sites (57.9%), with 58 of the 275 tested mosquitoes identified as kdr-positive. The lowest resistant allele frequency was 5% at the location Wechselburg (MW), and the highest values were 50% in Berlin (BE) and Frankfurt-Nordend (NOR). The ace-1 mutation was less abundant compared to the kdr mutation, consequently only five sites out of 38 were found positive (12% of sampling sites): Bad Camberg (LM_1), Glashütten (GLA), Frankfurt-Westend (WES), Frankfurt-Sossenheim (SOS) and Muggensturm (RA). Only six out of 283 tested mosquitoes were identified as ace-1 positive. The highest resistant allele frequency was 8% in Glashütten (GLA; Supplementary Table 2). In general, kdr mutations were widely distributed across the country, while ace-mutations were only present around the metropolitan area of Frankfurt (four out of five sampled locations) and Muggensturm (RA; one out of five sampled breeding sites; Figure 3). In the targeted GAM analysis, the three environmental variables (precipitation, mean temperature of the warmest quarter (BIO10), SMOD class) remained individually significant. However, model comparison showed that the best-supported GAM consisted of a linear effect model based on annual precipitation (BIO12; edf = 1, F = 12.65, p = 0.0013) together with sampling month and number of breeding sites as control variables. Neither the addition of the mean temperature of the warmest quarter (BIO10) nor SMOD class or an additional spatial smoothing improved model support. The best-supported model explained 39.5% of the deviance (adjusted R² = 0.249). Model diagnostics indicated approximately normal residuals, no evidence that the basis dimension was too low, and no problematic concurvity was detected, although residual variance was heteroscedastic (Figure 4, Supplement Tables 5-8).

**Figure 3:**
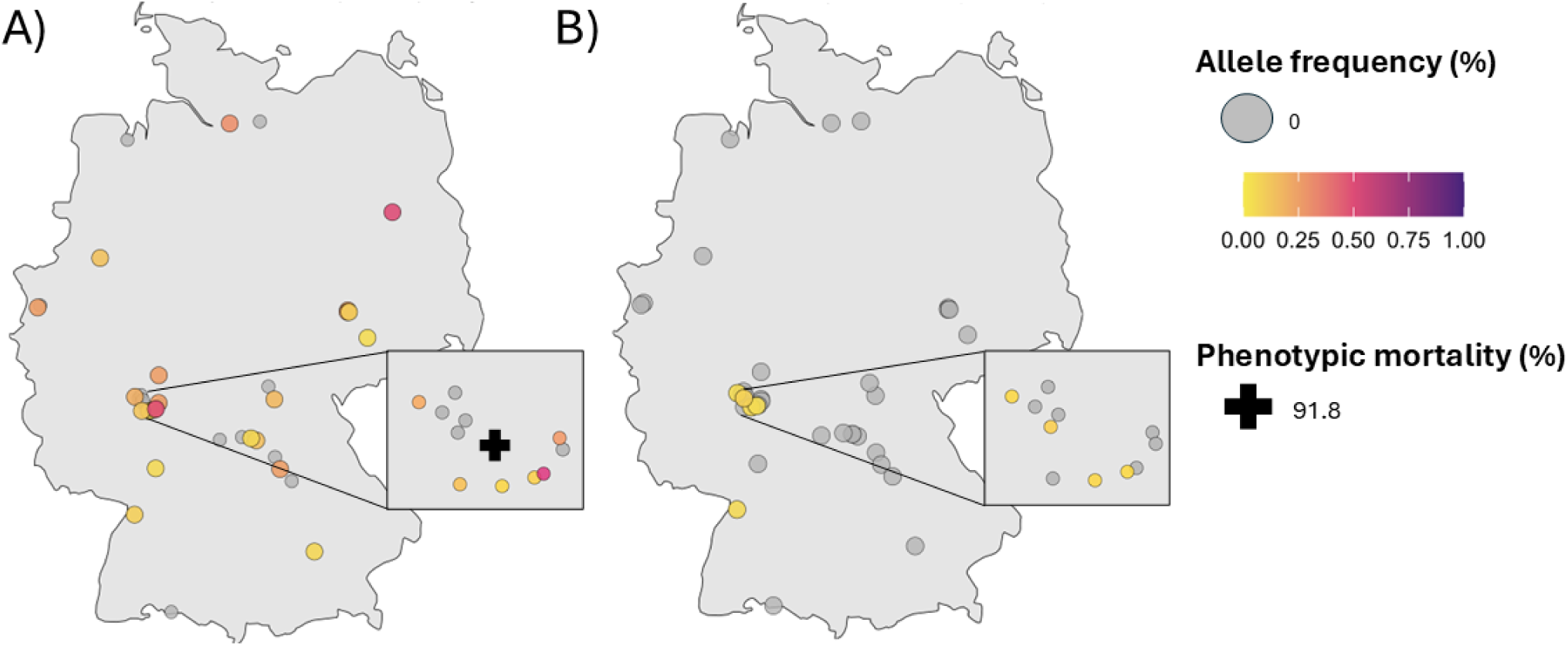
Spatial distribution of (A) kdr and (B) ace-1 resistance allele frequencies across Germany. The black cross indicates the origin of the *Cx. pipiens* f. *pipiens* population from Frankfurt used for phenotypic exposure to permethrin with the CDC bottle test. Not all sampling sites are represented, as kdr/ace-1 analysis could not be performed for individuals from all sites.

**Figure 4:**
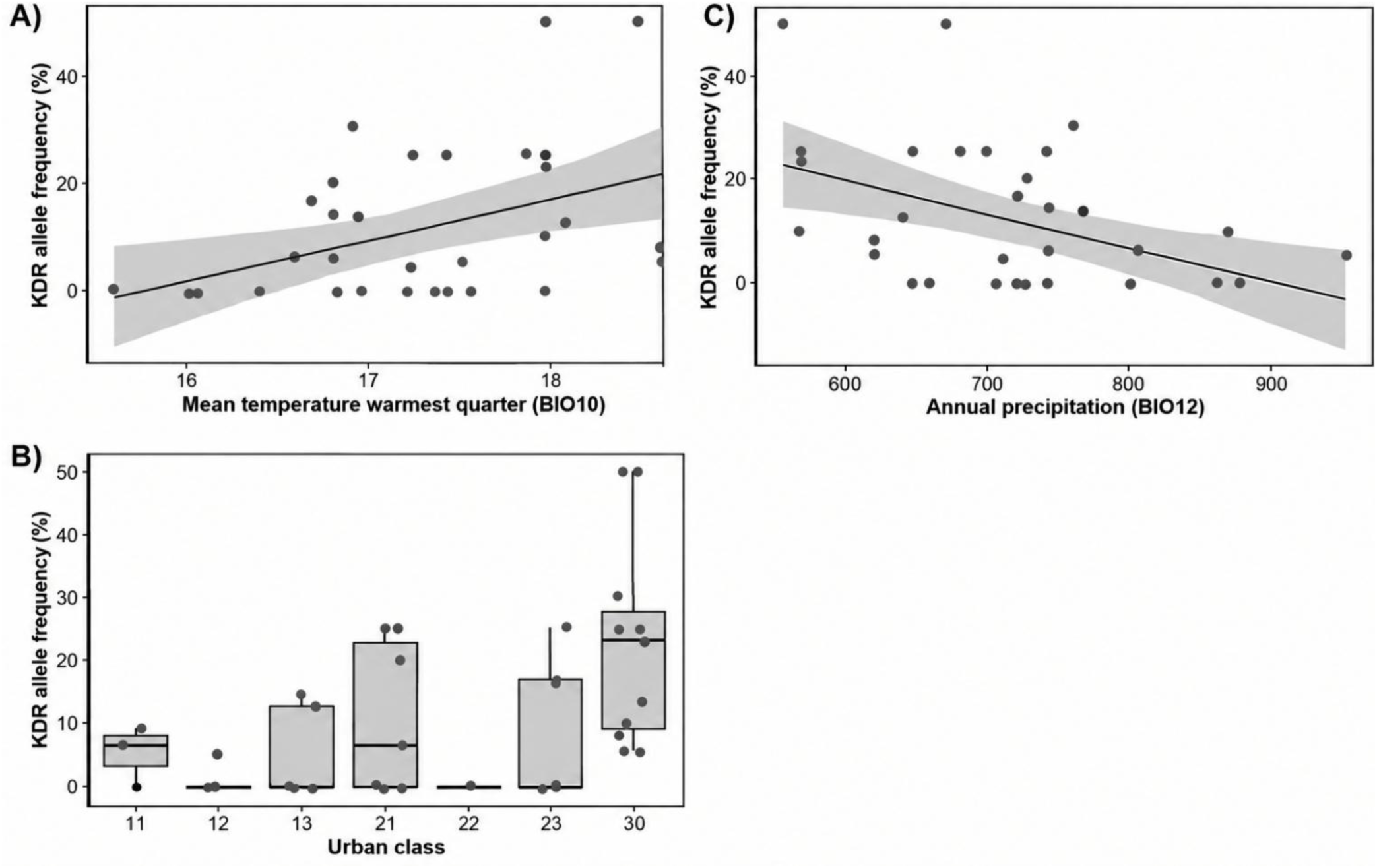
Relationships between environmental variables and kdr allele frequency across sampling sites in Germany. A) kdr allele frequency in relation to the mean temperature of the warmest quarter (BIO10) showing a linear response in the targeted GAM analysis; however, this predictor was not retained in the best-supported final GAM after accounting for control variables (sampling month and number of breeding sites sampled). B) kdr allele frequency in relation to SMOD class, showing significant differences among urban classes in the targeted analysis; however, this categorical predictor was not retained in the best-supported final GAM after accounting for the control variables. C) kdr allele frequency in relation to annual precipitation (BIO12), showing a significant linear relationship that remained robust in the final GAM after controlling for the control variables. Solid lines indicate fitted model predictions. Shaded areas represent 95% confidence intervals. Coloured dots are individual values per sampling location. The best-supported kdr GAM model included a linear effect of BIO12 and the control variables explaining 39.5% of deviance (adjusted R² = 0.249).

### Low haplotype diversity of *Culex pipiens* f. *pipiens*

The haplotype analysis using 84 COI sequences from 23 different sampling sites and nine federal states resulted in a network with five haplotypes (Figure 5). The network was dominated by one haplotype (H4) observed at all sampling sites. The haplotypes H1, H3, and H5 were only observed in one individual each. The second most common haplotype (H2) was found in seven populations (Figure 5). The populations from locations Hesse (FM), Saxony (RI) and Hesse (RB) show a higher genetic diversity compared to other populations, as they consist of three haplotypes. These three populations showed the second-highest haplotype diversity (Hd = 0.8), whereas the highest value was observed in the population Kempen (VIE) which contained only two sequences. Furthermore, the overall nucleotide diversity is rather low (π = 0.00427), as well as the haplotype diversity (H = 0.22088). AMOVA showed that 86% of genetic variation occurred within populations, and 14% of the genetic variation was attributable to differences between populations (Φ 0.135, df = 19) indicating weak population structure and low genetic differentiation. Notably, the two most frequent haplotypes (H2, H4) are separated by twelve mutations (Figure 5).

**Figure 5:**
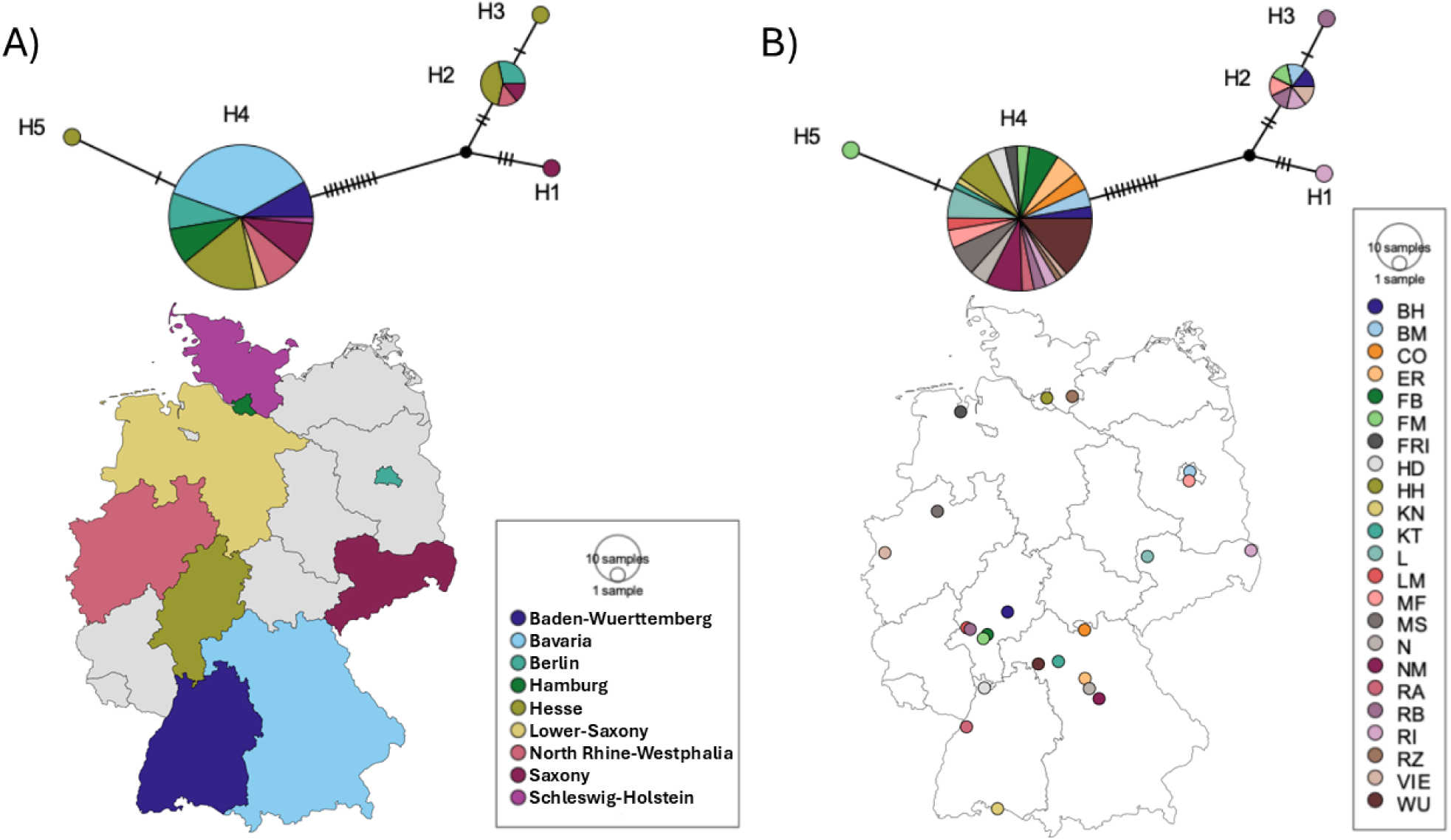
COI haplotype network of *Cx. pipiens* f. *pipiens* across Germany, using the TCS algorithm. Each circle depicts a unique haplotype, and the size is proportional to the number of sequences it represents. Nucleotide differences are show by the hatch marks across lines connecting the haplotypes. The dark unlabelled dots represent inferred ancestral nodes. A) Haplotype network per federal state (n=9) with colours corresponding to the legend. B) Haplotype network per sampling site (n=23) with colours corresponding to the legend.

Out of the 84 *Cx. pipiens* f. *pipiens* individuals subjected to COI barcoding for haplotype analysis, 61 were also analysed for molecular resistance at the kdr and ace-1 loci. However, no specific COI haplotype could be linked to resistance-associated mutations because 98% of these individuals belonged to haplotype H4 and 2% to haplotype H2 (Supplementary Table 9).

### Possible insecticide resistance of a population in the Frankfurt metropolitan area

The highest molecular kdr resistance was detected in locations in the metropolitan area of Frankfurt (Figure 3A). Thus, 127 females from Frankfurt were used for insecticide resistance phenotyping (Supplementary Table 4). The population was found to be possibly resistant to permethrin showing a mortality of 91.8% after the diagnostic time of 30 minutes (Figure 6). After 45 minutes, 100% mortality was observed in the exposed groups (Figure 6). Survival rates were 7.14% in the first and second test bottles, 11.11% in the third, and 8.33% in the fourth test bottle. Conversely, genotyping revealed no kdr resistance in the first test bottle (0%), but 14.28%, 5.55%, and 4.17% in the second, third, and fourth bottle, respectively. Accordingly, discrepancies were observed between phenotypic and molecular insecticide resistance results of the same mosquitoes tested (Figure 6).

**Figure 6:**
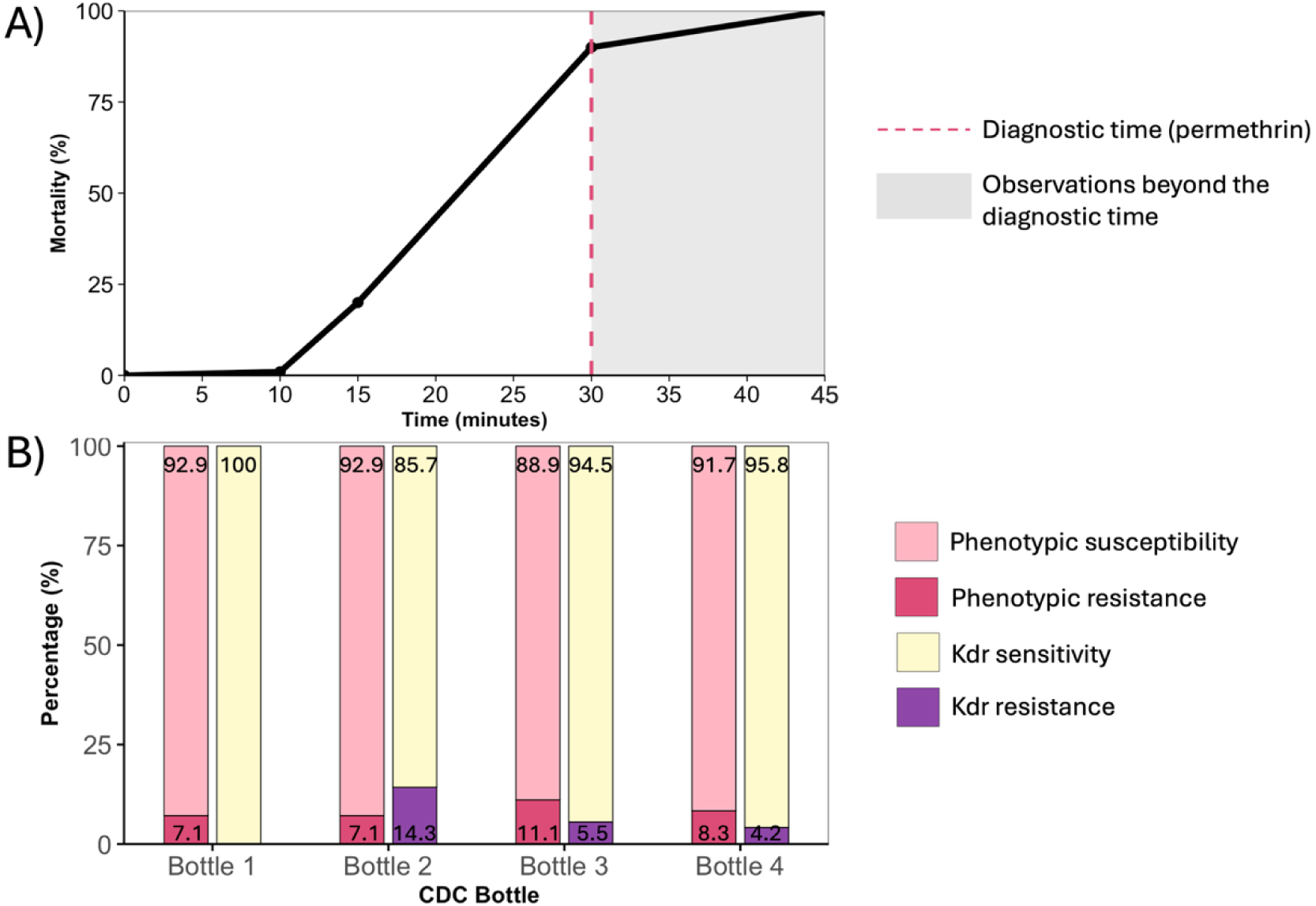
Genotypic and phenotypic insecticide resistance of a *Cx. pipiens* f. *pipiens* population from Frankfurt. A) Mortality results from the CDC bottle bioassay, representing the mean value of the four bottles. Dots represent the times at which mortality was counted, the black line shows the mortality curve. B) Molecular (kdr) and phenotypic (CDC bottle assay) insecticide resistance test results per CDC exposure bottle.

## Discussion

Molecular insecticide resistance across Germany, was found to be widespread and highly variable with particularly high levels observed in the metropolitan area Frankfurt am Main (Figure 3). Elevated kdr allele frequencies were correlated with annual precipitation (BIO12; Figure 3). COI haplotypes of *Cx*. *pipiens* f. *pipiens* could not be linked to a resistance genotype (Figure 5). The number of COI haplotypes per location of this study (one to five haplotypes per location) was comparable (one to six haplotypes per site) to the results by Werblow et al. (2014) although they did not differentiate bioforms (Figure 5; Werblow et al., 2014). To examine genotype–phenotype relationships in an area of high kdr resistance (Frankfurt am Main Nordend (NOR): 50%, Frankfurt am Main Botanical Gardens (WES): 8%), a *Cx*. *pipiens* f. *pipiens* population from the Frankfurt metropolitan area was phenotypically analysed and showed possible resistance to permethrin. However, genotype–phenotype discrepancies were observed, indicating that kdr mutations are only partly responsible for resistance in Germany (Figure 6).

### Environmental factors partially explain the distribution of molecular insecticide resistance

Consistent with our hypothesis, *Cx. pipiens* f*. pipiens* exhibited the L1014F and G119S resistance mutations in populations sampled in urban green spaces across the country (Figure 3). Resistance monitoring is of utmost importance, as WNV has been spreading in Germany since 2018, and authorities recommend the use of insecticides during outbreaks (Expertenkommission für Stechmücken am FLI, 2022). However, without knowledge of the resistance status, insecticide applications may be environmentally harmful and ineffective against mosquitoes. Accordingly, this study provides the first attempt of a broad-scale assessment of the molecular insecticide resistance status of German *Cx. pipiens* f*. pipiens* contributing to the risk assessment of vector control strategies. Kdr allele frequency was negatively associated with annual precipitation (BIO12; explaining 39.5 % of deviance). Under lower precipitation, reduced breeding-site availability may favour the spread of resistant genotypes by promoting smaller populations with reduced genetic variation (Weedall et al., 2020). Contrary to our hypothesis, kdr allele frequencies were not associated with temperature or urbanicity in the final model, but with precipitation-related environmental conditions. This potentially implies that broad-scale climatic conditions may currently be more important for structuring kdr variation than anthropogenic environmental gradients in German *Cx. pipiens* f. *pipiens* populations. Previous studies have linked insecticide resistance in mosquito populations to human activities such as urbanisation, domestic insecticide use, and agricultural pesticide application (Arich et al., 2024; Mouhamadou et al., 2019). However, such effects were not supported by our final model indicating that these drivers may be less important for *Cx. pipiens* f. *pipiens* at the spatial scale considered here. Another explanation is that the environmental factors at chosen sampling sites do not capture and resolve these human activities. Our sampling was restricted to urban green spaces, which may limit the generalisability of our findings. Broader sampling across cities and urbanisation gradients is needed to identify true urban effects. In addition to the high kdr frequencies, the ace-1 mutations were clustered in the metropolitan area of Frankfurt and were also present in Muggensturm (RA), which did not support our hypothesis of a broadly heterogeneous distribution across German populations. However, further studies based on broader sampling across a wider range of sites and land-use types, rather than focusing solely on urban green spaces, are needed to better disentangle the relative roles of climatic, ecological, and anthropogenic drivers of resistance patterns, especially as the kdr results should be interpreted with caution given the only moderate explanatory power of the best-supported model.

### Uniform genetic structure in *Culex pipiens* f*. pipiens*

COI haplotype analysis of *Cx. pipiens* f. *pipiens* revealed low genetic diversity, as hypothesised (Figure 5). Despite the low haplotype diversity, exceptions such as the Rietschen (RI), Reinborn (RB) and Frankfurt-Westend (WES; Supplementary Table 3) populations consisting of three different haplotypes, were present. The findings from Rietschen are consistent with those of Werblow et al. (2014), who reported the presence of more than three different haplotypes in this population (Werblow et al., 2014). On the other hand, our results showed no clear geographical pattern in the haplotype distribution of this species, indicating that the haplotypes are widespread. However, Werblow et al. (2014) could prove genetic variability and structure in eastern parts of Germany within *Cx. pipiens* s.l. compared to western populations (Werblow et al., 2014). The reason for this might be that they did not differentiate between the biotypes, while in this study only *Cx. pipiens* f. *pipiens* was investigated. Although nucleotide substitutions were detected among COI haplotypes in this study, these differences were not associated with geographic patterns suggesting that haplotype variation may be driven by non-spatial factors such as high gene flow. This lack of geographic structuring in *Cx. pipiens* f. *pipiens* further indicates that COI is too conservative to resolve fine-scale population differentiation. In addition, given the limited COI haplotype diversity, no association between COI haplotypes and resistance-associated mutations could be detected.

### Weak phenotype–genotype link in insecticide resistance

Phenotypic testing of a *Cx. pipiens* f. *pipiens* population from Frankfurt indicated possible resistance to permethrin (91.8%). This might indicate early stages of resistance development in Germany. Other European countries are already reporting resistance to permethrin (Vereecken et al., 2022). Our findings of possible phenotypic resistance in *Cx. pipiens* f*. pipiens* from Frankfurt am Main, is in line with the recent study, which topically exposed *Cx. pipiens* f. *molestus* individuals from South-Baden and *Cx. pipiens* s.l. individuals from Heidelberg to permethrin and found potential resistance (Portwood et al., 2025). Eventually, Portwood et al. (2025) concluded a potential trend towards permethrin-resistance in German *Cx. pipiens* populations. However, the results are difficult to compare because different resistance assays were used: The CDC bottle bioassay is a standard method, whereas the topical exposure test applied by Portwood et al. (2025) is a more recent approach, which complicates direct comparisons (Portwood, et al., 2025). However, both *Cx. pipiens* biotypes sampled in different cities in Germany express possible resistance to permethrin. In the neighbouring country Belgium, a population of *Cx. pipiens* f*. molestus* was found resistant to permethrin with a mortality of 78% using the WHO susceptibility assay (Vereecken et al., 2022). A possible explanation could be that the Belgian population consisted of *Cx. pipiens* f*. molestus*, a biotype known to be active throughout the whole year and preferring to bite humans over birds, thus living close to humans and being potentially more exposed to insecticides used in households (Gray et al., 2018). Moreover, our study relied on the use of the CDC bottle assay rather than the WHO test tubes for assessing insecticide susceptibility. Although both assays are widely used (ECDC, 2023), differences in diagnostic doses, exposure times and handling of mosquitoes can influence mortality outcomes and comparison with Vereecken et al. (2022).

In the tested Frankfurt population, no genotype-phenotype link was detected (Figure 6). Although a few individuals exhibited phenotypic resistance, the presence of the kdr mutation did not consistently correspond to the observed resistance patterns across the CDC bottles, thereby not supporting our hypothesis that populations with elevated molecular resistance markers exhibit corresponding phenotypic resistance and indicating that kdr-mediated resistance alone cannot explain the phenotypic outcomes. Further research is required to better understand the factors shaping the insecticide resistance in *Cx. pipiens* f. *pipiens* in Germany. The resistance mechanism of the kdr mutation is a modified binding site of the pyrethroid insecticide (Brengues et al., 2003). Other resistance mechanisms such like adapted detoxification processes or the modulation of the mosquito cuticle (thickening, insecticide degradation at target sites or modulation of binding receptors) may play an additional role (Ingham et al., 2023). Thus, genotype–phenotype relationships remain inconclusive, but the broad scale molecular survey provides valuable clues about resistance status and serves as a foundation for future investigations into insecticide resistance mechanisms in German *Cx. pipiens*.

## Limitation of the study

A potential limitation of the low COI diversity in *Cx. pipiens* f. *pipiens* individuals may partly reflect a sampling limitation, as sampling was restricted to urban green spaces located near human settlements. Therefore, individuals were collected in similar habitats and in a limited number of breeding grounds (Figure 2A), which may lead to reduced haplotype diversity because of similar ecological niches and familial relatedness. However, we coped for that by randomly selecting individuals for sequencing (Supplementary Table 1). Our sampling was restricted to urban green spaces, and a large proportion of the sampling sites were in and around Frankfurt/Hesse, which may limit the generalisability of our findings. Similar high resistance patterns as in Frankfurt may nevertheless occur in other highly urbanised regions, such as the capital Berlin. As similar habitats were sampled in general (urban green spaces), broader sampling across land use-gradients is needed to better assess the effects of urbanisation on molecular resistance.

## Conclusion

This is the first study that presents comprehensive data on molecular insecticide resistance levels in *Cx. pipiens* f. *pipiens* across Germany. Molecular analyses identified the presence of insecticide resistance-associated mutations across the country. Kdr allele frequency was linked to precipitation suggesting that climatic conditions may be more important than anthropogenic factors (urbanicity) in shaping resistance patterns, although the observed pattern was only moderately supported by the model (Figure 4). In addition to molecular resistance testing, phenotypic insecticide resistance was assessed in Germany using a standardised CDC bioassay, revealing emerging permethrin resistance in a *Cx. pipiens* f. *pipiens* population from Frankfurt am Main, a metropolitan region characterised by high levels of molecular resistance. However, no strong resistance genotype-phenotype link was detected suggesting that kdr-mediated resistance alone cannot explain the phenotypic outcomes. COI haplotype analyses revealed low genetic variability between *Cx. pipiens* f. *pipiens* populations, with one dominant haplotype, preventing an assessment of associations with resistance patterns (Figure 5). Overall, our findings provide valuable baseline data for future insecticide resistance monitoring, while comprehensive nationwide surveillance studies in Germany, encompassing underrepresented regions and a broad range of habitat types, are needed to assess patterns of insecticide resistance across the country, to improve risk assessment and inform evidence-based vector control strategies.

## Acknowledgements

The authors wish to thank Henrik Hartke, Jessica Neid, Johanna Bock, Juliane Hartke, Marina Psalti, Markus Braun, Pia Guckert, Simon Prudlo, Susanne Hartke, Timo Pampuch, Winfried Bock, Lotta Müller and Florian Giertz for the sampling efforts during the sampling campaign in 2023. Furthermore, the authors would like to thank the Botanical Gardens at Frankfurt am Main, Germany, for trap maintenance and granting access to the site for sample collection.

## Author contributions

IMK, FR and RM conceptualised the study. RM and DG supplied facilities and equipment. IMK and FR were responsible for the mosquito larval collection with help of AM. EV performed all laboratory work. SV helped with the insecticide resistance phenotyping. CJ helped with molecular work. EV and IB analysed the COI sequences. EV generated, analysed, and visualised the data with supervision of IMK. The work was part of EV’s master thesis which was supervised by IMK and RM. EV wrote the original manuscript draft. All authors reviewed and edited the draft. All authors read and approved of the final manuscript.

## Supplementary information

The following supplementary material is available.

Additional file 1:

*Supplementary Figure 1: Mosquito sampling locations across Germany*.

*Supplementary Figure 2: Climatic conditions at sampling sites in 2023*.

*Supplementary Figure 3: Nineteen extracted bioclimatic variables (Karger et al., 2017)*.

*Supplementary Figure 4: Urban land use classes at sampling sites*.

*Supplementary Figure 5: Species co-occurrence per breeding site. Locations of breeding sites are given in Supplementary Table 1*.

*Supplementary Table 1: Characteristics of sampling success per site with number of individuals successfully identified, number of breeding habitats and amount of different species identified*.

*Supplementary Table 2: Culex pipiens f. pipiens individuals that were subjected to kdr and ace-1 analysis*.

*Supplementary Table 1: Cx. pipiens f. pipiens samples used for COI haplotype analysis*

*Supplementary Table 4: Mortality results from the CDC bottle bioassay of Cx. pipiens f. pipiens. PR = possible resistance*.

*Supplementary Table 2: Targeted GAM screening results*.

*Supplementary Table 3: The coefficients and smooth terms of the best-supported final GAM*

*Supplementary Table 4: Comparison of the candidate GAMs using AIC and AICc*.

*Supplementary Table 5: Diagnostic statistics for the best-supported GAMs*.

Additional file 2:

*Supplementary Table 9: Results of the molecular species identification, insecticide resistance mutations and haplotype analysis for each sampled mosquito*.

## Data availability

All data generated and analysed during this study are included within this article and its additional files. All COI sequences are available in GenBank under accession numbers PZ861051 - PZ861134.

