## Supplementary file 1 for "Insecticide resistance of *Culex pipiens* f. *pipiens* is geographically widespread in Germany and displays an unmatched genotypic and phenotypic resistance state"

**Supplementary Information**

*Supplementary Figures*

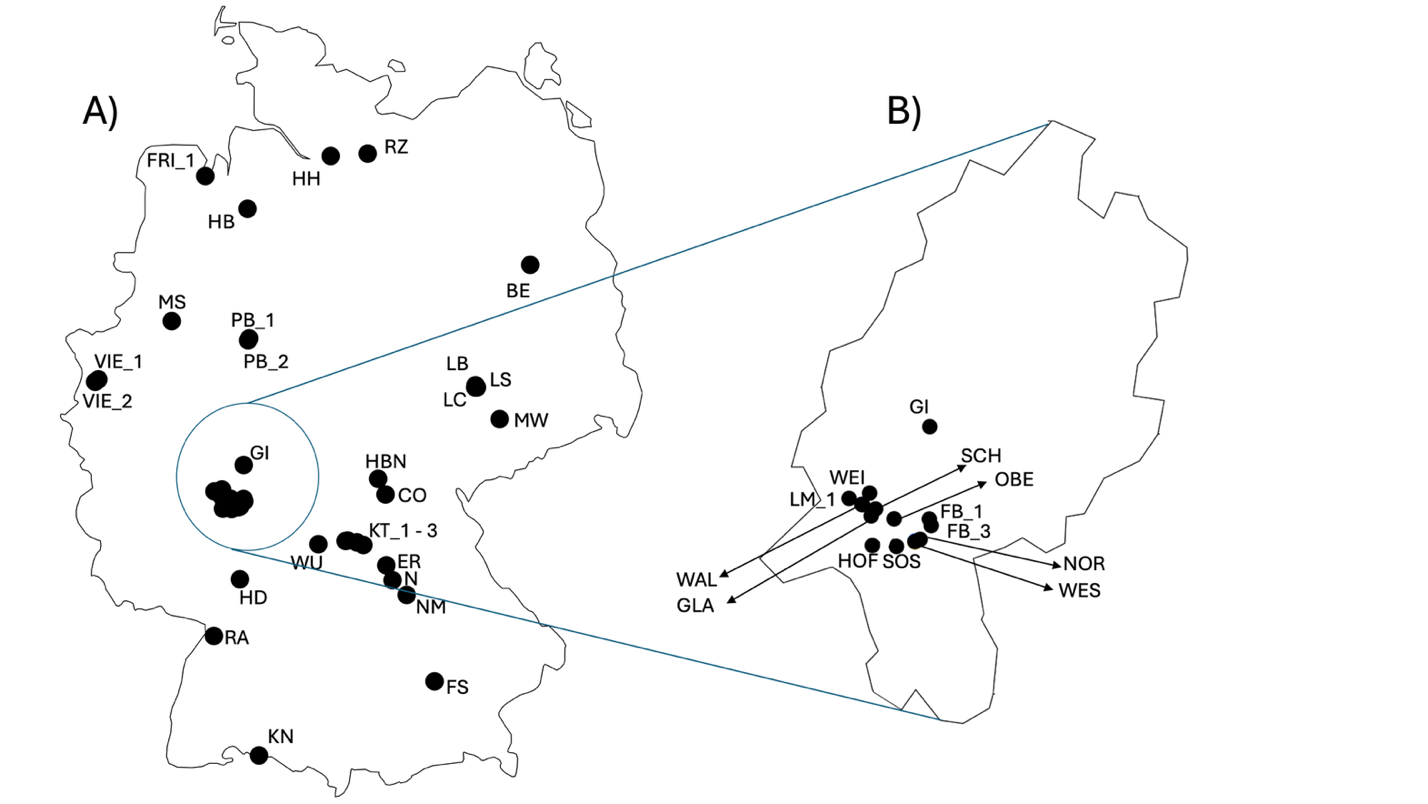

Supplementary Figure 1: Mosquito sampling locations across Germany. A) Overview of all the sampling sites (details in Supplementary Table 1). B) Focus on the federal state of Hesse. The sampling sites are indicated with their abbreviation which is equivalent to other further Figures showing results. Full names of cities are present in Supplementary Table 1.

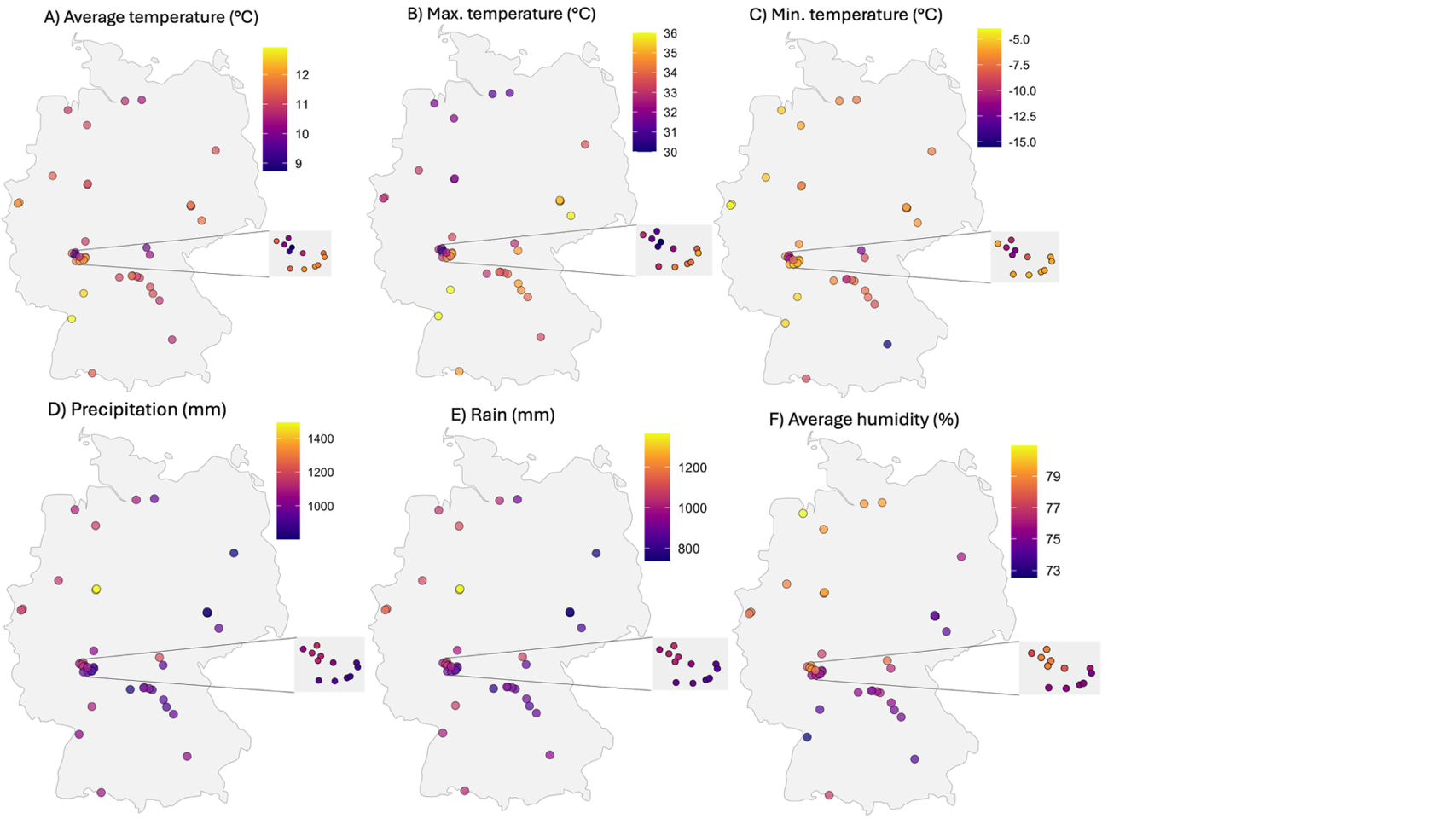

Supplementary Figure 2: Climatic conditions across sampling sites in 2023. A) Average annual air temperature (°C). B) Maximum temperature (°C). C) Minimum temperature (°C). D) Sum of precipitation (mm). E) Sum of rain (mm). F) Average humidity (%). Extracted from the Historical Weather API.

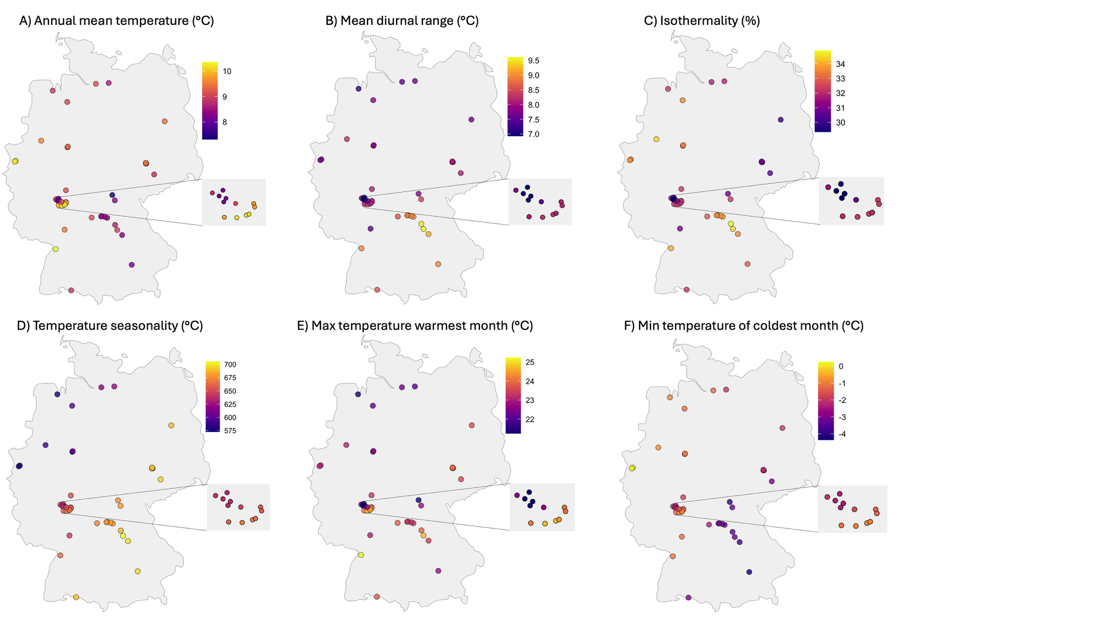

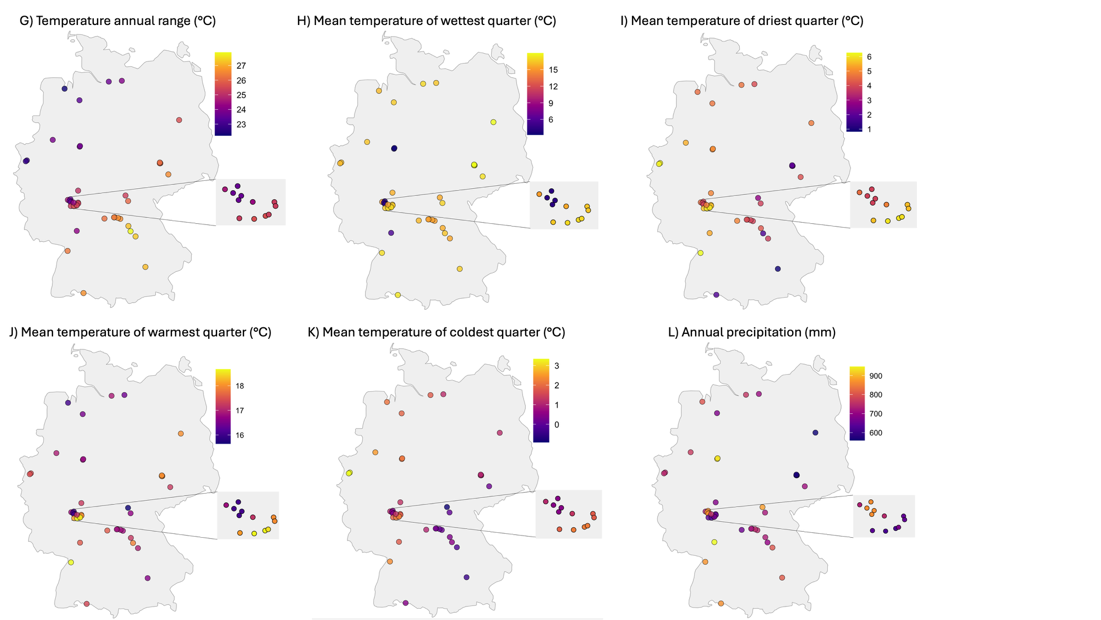

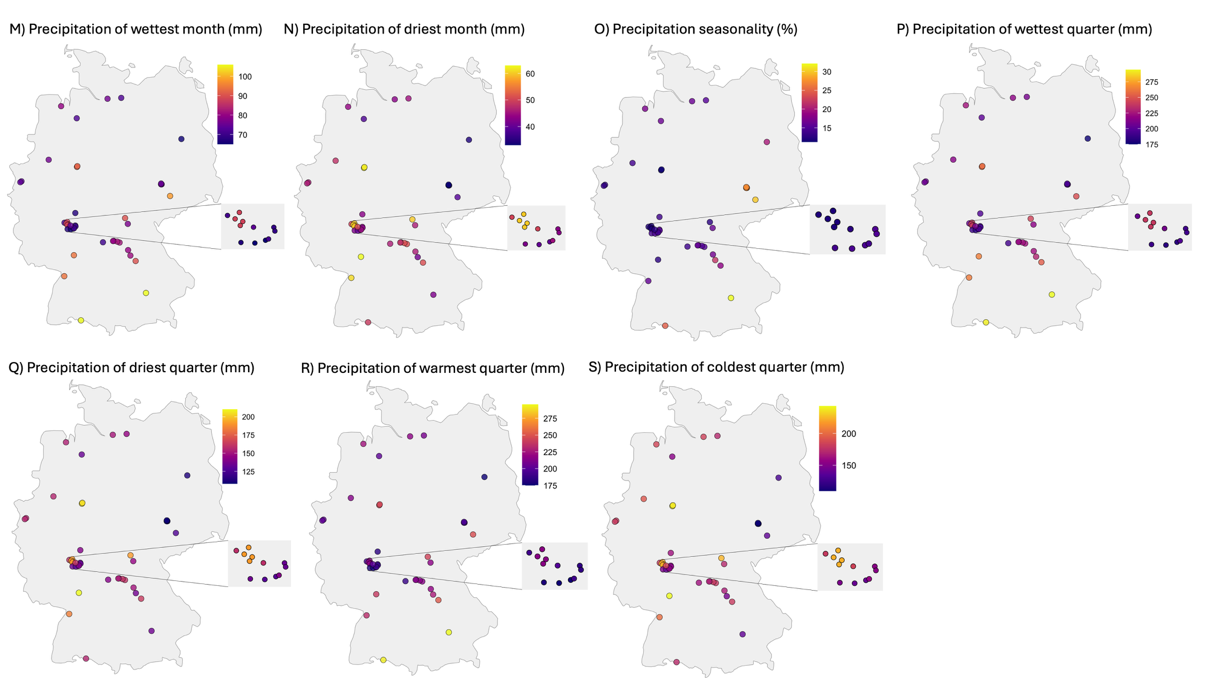

Supplementary Figure 3: Nineteen extracted Bioclimatic variables (Karger et al., 2017).

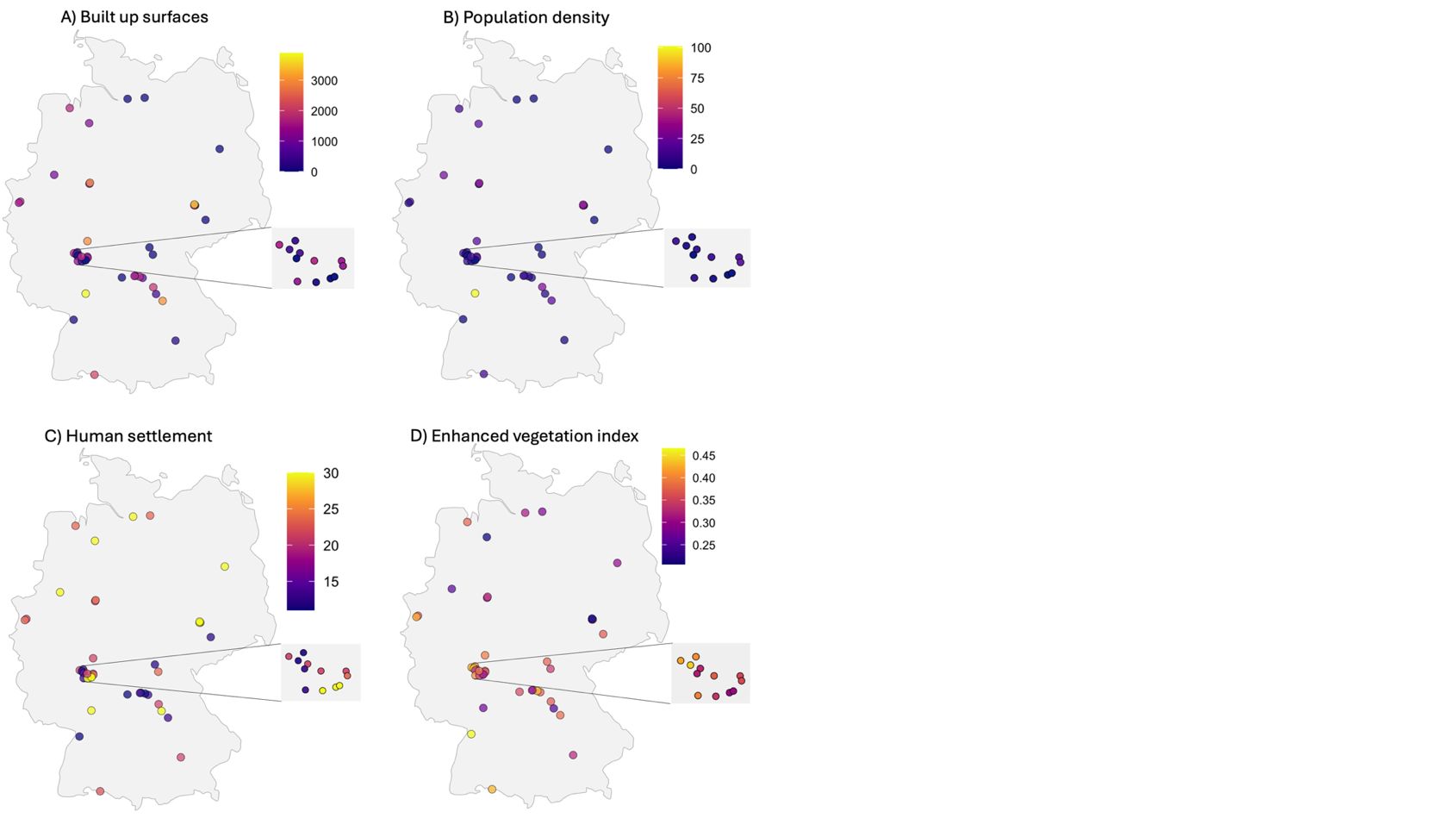

Supplementary Figure 4: Urban land use patterns across sampling sites. A) Built-up surfaces per 10,000m^2^. B) Population density (people/10,000m^2^). C) Human settlement classes. Extracted from the THS-built dataset of the year 2020 (Ehrlich et al., 2018). D) The enhanced vegetation index, extracted from the NASA of the year 2023.

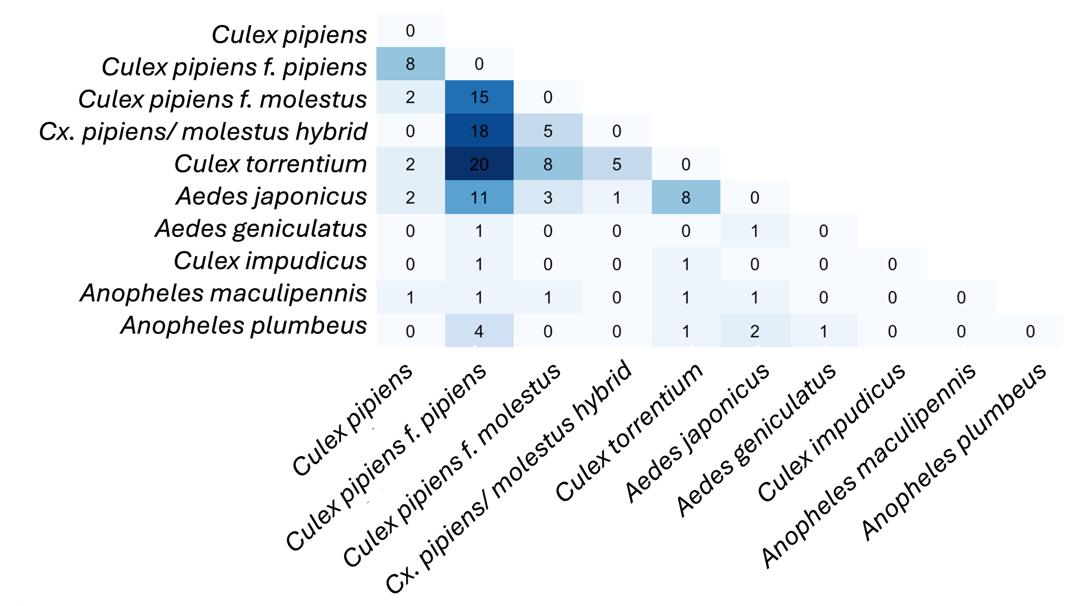

Supplementary Figure 5: Number of habitats where species pairs co-occurs.

Supplementary Tables

Supplementary Table 1: overview of sampling sites with number of individuals successfully identified to species level, number of sampled breeding habitats and number of different species identified per sampling site.

| **Site** | **City (Federal state)** | **Latitude, Longitude** | **Number of individuals identified** | **Number of breeding habitats** | **Number of species identified** |
| --- | --- | --- | --- | --- | --- |
| BA_1 | Schlüsselfeld (Bavaria) | 49.75606, 10.62595 | 23 | 2 | 2 |
| BE | Berlin (Berlin) | 52.513784, 13.252367 | 9 | 1 | 3 |
| CO | Coburg (Bavaria) | 50.25341, 10.97477 | 35 | 3 | 2 |
| ER | Erlangen (Bavaria) | 49.55588, 10.98726 | 21 | 3 | 3 |
| FB_1 | Bad Vilbel (Hesse) | 50.20947, 8.73995 | 13 | 2 | 2 |
| FB_3 | Bad Vilbel (Hesse) | 50.18592, 8.75060 | 37 | 1 | 3 |
| FRI_1 | Varel (Lower Saxony) | 53.38779, 8.13890 | 22 | 2 | 2 |
| FS | Freising (Bavaria) | 48.413965, 11.746977 | 25 | 2 | 3 |
| GI | Pohlheim (Hesse) | 50.543048, 8.742562 | 18 | 2 | 5 |
| HB | Bremen (Bremen) | 53.065722, 8.803305 | 10 | 2 | 1 |
| HBN | Eisfeld (Thuringia) | 50.40770, 10.85917 | 17 | 1 | 3 |
| HD | Heidelberg (Baden-Wuerttemberg) | 49.419184, 8.682792 | 54 | 2 | 3 |
| HH | Hamburg (Hamburg) | 53.585000, 10.112111 | 17 | 1 | 1 |
| KN | Gaienhofen (Baden-Wuerttemberg) | 47.68429, 8.98365 | 7 | 2 | 4 |
| KT_1 | Wiesentheid (Bavaria) | 49.80001, 10.37773 | 17 | 2 | 1 |
| KT_2 | Geiselwind (Bavaria) | 49.78248, 10.52311 | 34 | 5 | 5 |
| KT_3 | Wiesentheid (Bavaria) | 49.79620, 10.34708 | 5 | 2 | 2 |
| LM_1 | Bad Camberg (Hesse) | 50.284043, 8.283191 | 35 | 1 | 4 |
| MS | Münster (North Rhine-Westphalia)  ) | 51.960208, 7.610165 | 57 | 1 | 3 |
| MW | Wechselburg (Saxony) | 50.995947, 12.771589 | 44 | 3 | 3 |
| N | Neurenberg (Bavaria) | 49.40993, 11.08463 | 43 | 2 | 2 |
| NM | Pyrbaum (Bavaria) | 49.26457, 11.30795 | 16 | 2 | 3 |
| PB_1 | Bad Lippspringe (North Rhine-Westphalia)  ) | 51.77041, 8.81240 | 3 | 1 | 2 |
| PB_2 | Bad Lippspringe (North Rhine-Westphalia)  ) | 51.79029, 8.82912 | 9 | 2 | 5 |
| RA | Muggensturm (Baden-Wuerttemberg) | 48.860838, 8.273359 | 35 | 1 | 1 |
| RZ | Mölln (Schleswig-Holstein) | 53.608667, 10.694611 | 13 | 3 | 2 |
| VIE_1 | Kempen (North Rhine-Westphalia)  ) | 51.386675, 6.455026 | 7 | 1 | 1 |
| VIE_2 | Kempen (North Rhine-Westphalia)  ) | 51.362588, 6.405421 | 33 | 2 | 2 |
| WÜ | Würzburg (Bavaria) | 49.76395, 9.91819 | 16 | 2 | 1 |
| WEI | Weilrod-Riedelbach (Hesse) | 50.303118, 8.400111 | 21 | 1 | 3 |
| SCH | Schmitten-Oberreifenberg (Hesse) | 50.245881, 8.433824 | 41 | 1 | 2 |
| WAL | Waldems-Reinborn (Hesse) | 50.262412, 8.358149 | 43 | 1 | 3 |
| GLA | Glashütten (Hesse) | 50.220793, 8.408946 | 59 | 1 | 3 |
| HOF | Hofheim-Lorsbach (Hesse) | 50.114053, 8.415923 | 55 | 1 | 4 |
| WES | Frankfurt-Westend (Hesse) | 50.128030, 8.658117 | 32 | 2 | 4 |
| SOS | Frankfurt-Sossenheim (Hesse) | 50.110439, 8.553031 | 18 | 2 | 3 |
| NOR | Frankfurt-Nordend (Hesse) | 50.135983, 8.687971 | 21 | 1 | 2 |
| LS | Leipzig (Saxony) | 51.307237, 12.414632 | 82 | 1 | 3 |
| LC | Leipzig (Saxony) | 51.306477, 12.387084 | 66 | 2 | 3 |
| LB | Leipzig (Saxony) | 51.327956, 12.391006 | 46 | 3 | 2 |
| OBE | Oberursel (Hesse) | 50.210142, 8.539765 | 38 | 1 | 2 |
| BH | Breungeshain ( Hesse) | 50.511505, 9.21294662 | 3 | 1 | 1 |
| RI | Rietschen (Saxony) | 51.3951637, 14.784715 | 4 | 1 | 1 |
| RB | Reinborn (Hesse) | 50.2625571, 8.35807375 | 4 | 1 | 1 |
| MF | Berlin-Marienfelde (Berlin) | 52.39626, 13.36647 | 4 | 1 | 1 |
| BM | Berlin-Mitte (Berlin) | 52.5280984, 13.3842021 | 4 | 1 | 1 |
| FM | Frankfurt-Westend (Hesse) | 50.1290507, 8.65756914 | 4 | 1 | 1 |

Supplement Table 2: Cx. pipiens f. pipiens individuals sampled in Germany that were subjected to kdr and ace-1 PCR. Individuals that failed the PCR were excluded from further analyses. RS = heterozygous resistant; SS = homozygous sensitive; RR = homozygous resistant.

| **Larval ID** | **Site** | **Habitat** | **Sampling month** | **Kdr PCR** | **Ace-1 PCR** |
| --- | --- | --- | --- | --- | --- |
| HE_FB_09_02 | FB_1 | garden | July | failed | SS |
| HE_FB_09_05 | FB_1 | garden | July | RS | SS |
| HE_FB_09_06 | FB_1 | garden | July | RS | SS |
| HE_FB_09_09 | FB_1 | garden | July | SS | SS |
| HE_FB_09_16 | FB_1 | garden | July | SS | SS |
| HE_FB_33_02 | FB_3 | graveyard | July | SS | SS |
| HE_FB_33_07 | FB_3 | graveyard | July | SS | SS |
| HE_FB_33_10 | FB_3 | graveyard | July | SS | SS |
| HE_FB_33_15 | FB_3 | graveyard | July | SS | SS |
| HE_FB_33_19 | FB_3 | graveyard | July | SS | SS |
| NI_FRI_46_01 | FRI_1 | graveyard | July | SS | SS |
| NI_FRI_46_15 | FRI_1 | graveyard | July | SS | SS |
| NRW_PB_50_15 | PB_1 | garden | July | failed | failed |
| BW_KN_52_09 | KN | village square | July | SS | SS |
| BW_KN_52_10 | KN | village square | July | SS | SS |
| BW_KN_52_11 | KN | village square | July | SS | SS |
| BY_KT_70_11 | KT_1 | graveyard | August | SS | failed |
| BY_KT_70_16 | KT_1 | graveyard | August | SS | SS |
| BY_KT_71_08 | KT_2 | garden | August | SS | SS |
| BY_KT_71_13 | KT_2 | garden | August | SS | SS |
| BY_KT_72_02 | KT_2 | garden | August | failed | SS |
| BY_KT_72_16 | KT_2 | garden | August | RS | SS |
| BY_N_73_02 | N | graveyard | August | failed | failed |
| BY_N_73_11 | N | graveyard | August | failed | failed |
| BY_N_73_14 | N | graveyard | August | failed | failed |
| BY_N_74_06 | N | graveyard | August | SS | SS |
| BY_N_74_11 | N | graveyard | August | SS | SS |
| BY_N_74_18 | N | graveyard | August | failed | failed |
| BY_N_75_06 | N | graveyard | August | RS | SS |
| BY_N_75_09 | N | graveyard | August | RS | SS |
| BY_N_75_15 | N | graveyard | August | RS | SS |
| BY_N_75_17 | N | graveyard | August | RS | SS |
| BY_NM_76_01 | NM | graveyard | August | SS | SS |
| BY_NM_76_05 | NM | graveyard | August | SS | SS |
| BY_NM_76_06 | NM | graveyard | August | SS | SS |
| BY_NM_76_07 | NM | graveyard | August | SS | SS |
| BY_NM_76_08 | NM | graveyard | August | SS | SS |
| BY_NM_76_11 | NM | graveyard | August | SS | SS |
| BY_NM_76_13 | NM | graveyard | August | SS | SS |
| BY_WÜ_78_01 | WÜ | graveyard | August | SS | SS |
| BY_WÜ_78_02 | WÜ | graveyard | August | SS | SS |
| BY_WÜ_78_03 | WÜ | graveyard | August | SS | SS |
| BY_WÜ_78_04 | WÜ | graveyard | August | SS | SS |
| BY_WÜ_78_08 | WÜ | graveyard | August | SS | SS |
| BY_WÜ_78_11 | WÜ | graveyard | August | SS | SS |
| BY_WÜ_78_12 | WÜ | graveyard | August | SS | SS |
| BY_WÜ_78_13 | WÜ | graveyard | August | SS | SS |
| BY_WÜ_78_15 | WÜ | graveyard | August | SS | SS |
| BY_WÜ_78_16 | WÜ | graveyard | August | SS | SS |
| BY_CO_86_03 | CO | graveyard | August | SS | SS |
| BY_CO_86_07 | CO | graveyard | August | SS | SS |
| BY_CO_86_10 | CO | graveyard | August | SS | SS |
| BY_CO_86_17 | CO | graveyard | August | SS | SS |
| BY_CO_86_21 | CO | graveyard | August | failed | SS |
| BY_CO_87_03 | CO | graveyard | August | SS | SS |
| BY_CO_87_07 | CO | graveyard | August | RR | SS |
| BY_CO_87_10 | CO | graveyard | August | failed | SS |
| BY_CO_87_14 | CO | graveyard | August | SS | SS |
| BY_CO_87_20 | CO | graveyard | August | SS | SS |
| TH_HBN_88_08 | HBN | graveyard | August | SS | SS |
| BY_KT_90_05 | KT_3 | graveyard | August | SS | SS |
| BY_KT_90_09 | KT_3 | graveyard | August | SS | SS |
| BY_KT_91_19 | KT_2 | garden | August | SS | SS |
| BY_BA_92_03 | BA_1 | graveyard | August | SS | SS |
| BY_BA_92_08 | BA_1 | graveyard | August | SS | SS |
| BY_BA_92_10 | BA_1 | graveyard | August | RS | SS |
| BY_BA_92_12 | BA_1 | graveyard | August | failed | SS |
| BY_BA_92_18 | BA_1 | graveyard | August | SS | SS |
| BY_BA_94_01 | BA_1 | graveyard | August | failed | SS |
| BY_BA_94_05 | BA_1 | graveyard | August | failed | failed |
| BY_BA_94_07 | BA_1 | graveyard | August | failed | SS |
| BY_BA_94_09 | BA_1 | graveyard | August | SS | SS |
| BY_BA_94_18 | BA_1 | graveyard | August | SS | SS |
| BY_ER_99_15 | ER | graveyard | August | failed | failed |
| BY_ER_99_20 | ER | graveyard | August | failed | failed |
| BY_ER_101_05 | ER | graveyard | August | failed | failed |
| BY_ER_101_11 | ER | graveyard | August | SS | SS |
| BY_ER_101_14 | ER | graveyard | August | SS | SS |
| BY_ER_101_17 | ER | graveyard | August | SS | SS |
| BY_ER_101_19 | ER | graveyard | August | SS | SS |
| SN_MW_104_02 | MW | garden | August | failed | failed |
| SN_MW_104_08 | MW | garden | August | failed | SS |
| SN_MW_104_14 | MW | garden | August | SS | SS |
| SN_MW_104_16 | MW | garden | August | failed | SS |
| SN_MW_105_07 | MW | garden | August | failed | SS |
| SN_MW_105_12 | MW | garden | August | SS | SS |
| SN_MW_105_16 | MW | garden | August | SS | SS |
| SN_MW_106_10 | MW | garden | August | SS | SS |
| SN_MW_106_14 | MW | garden | August | SS | SS |
| SN_MW_106_16 | MW | garden | August | SS | failed |
| SN_L_108_06 | LS | graveyard | August | failed | failed |
| SN_L_108_14 | LS | graveyard | August | SS | SS |
| SN_L_109_07 | LS | graveyard | August | failed | failed |
| SN_L_109_13 | LS | graveyard | August | SS | SS |
| SN_L_110_15 | LS | graveyard | August | SS | SS |
| SN_L_110_19 | LS | graveyard | August | SS | SS |
| SN_L_111_10 | LS | graveyard | August | SS | SS |
| SN_L_111_12 | LS | graveyard | August | SS | SS |
| SN_L_112_03 | LS | graveyard | August | RS | SS |
| SN_L_112_06 | LS | graveyard | August | RS | SS |
| SN_L_113_07 | LC | graveyard | August | SS | SS |
| SN_L_113_09 | LC | graveyard | August | RS | SS |
| SN_L_113_16 | LC | graveyard | August | SS | SS |
| SN_L_114_05 | LC | graveyard | August | SS | SS |
| SN_L_114_17 | LC | graveyard | August | SS | SS |
| SN_L_115_08 | LC | graveyard | August | SS | SS |
| SN_L_115_18 | LC | graveyard | August | failed | SS |
| SN_L_117_04 | LC | graveyard | August | RS | SS |
| SN_L_117_11 | LC | graveyard | August | RS | SS |
| SN_L_117_18 | LC | graveyard | August | failed | failed |
| SN_L_118_06 | LB | garden | August | SS | SS |
| SN_L_118_08 | LB | garden | August | RS | SS |
| SN_L_118_10 | LB | garden | August | RS | SS |
| SN_L_119_12 | LB | garden | August | RS | SS |
| SN_L_119_18 | LB | garden | August | RR | SS |
| SN_L_120_16 | LB | garden | August | RS | failed |
| SN_L_120_18 | LB | garden | August | SS | SS |
| SN_L_121_08 | LB | garden | August | SS | failed |
| SN_L_121_14 | LB | garden | August | SS | SS |
| SN_L_121_18 | LB | garden | August | SS | failed |
| HE_GI_122_03 | GI | garden | September | RS | SS |
| HE_GI_122_20 | GI | garden | September | SS | SS |
| BY_FS_129_04 | FS | graveyard | September | failed | SS |
| BY_FS_129_06 | FS | graveyard | September | SS | SS |
| BY_FS_129_07 | FS | graveyard | September | SS | SS |
| BY_FS_129_09 | FS | graveyard | September | failed | failed |
| BY_FS_129_11 | FS | graveyard | September | SS | SS |
| BY_FS_129_14 | FS | graveyard | September | SS | SS |
| BY_FS_129_15 | FS | graveyard | September | SS | SS |
| BY_FS_129_16 | FS | graveyard | September | RS | SS |
| BY_FS_129_17 | FS | graveyard | September | SS | SS |
| BY_FS_129_19 | FS | graveyard | September | failed | SS |
| BW_RA_130_04 | RA | graveyard | October | SS | failed |
| BW_RA_130_16 | RA | graveyard | October | SS | SS |
| BW_RA_130_18 | RA | graveyard | October | RS | SS |
| BW_RA_131_04 | RA | graveyard | October | SS | SS |
| BW_RA_131_12 | RA | graveyard | October | RS | failed |
| BW_RA_131_18 | RA | graveyard | October | failed | failed |
| BW_RA_133_08 | RA | graveyard | October | SS | SS |
| BW_RA_133_10 | RA | graveyard | October | SS | SS |
| BW_RA_133_11 | RA | graveyard | October | SS | SS |
| BW_RA_133_13 | RA | graveyard | October | SS | SS |
| NRW_MS_134_04 | MS | graveyard | September | RS | SS |
| NRW_MS_134_15 | MS | graveyard | September | RS | SS |
| NRW_MS_134_20 | MS | graveyard | September | failed | failed |
| NRW_MS_135_03 | MS | graveyard | September | SS | SS |
| NRW_MS_135_08 | MS | graveyard | September | SS | SS |
| NRW_MS_135_14 | MS | graveyard | September | SS | SS |
| NRW_MS_135_21 | MS | graveyard | September | SS | SS |
| NRW_MS_136_06 | MS | graveyard | September | SS | SS |
| NRW_MS_136_10 | MS | graveyard | September | RS | SS |
| NRW_MS_136_16 | MS | graveyard | September | failed | SS |
| BE_BE_138_20 | BE | graveyard | September | RS | failed |
| HH_HH_01_02 | HH | graveyard | September | SS | SS |
| HH_HH_01_06 | HH | graveyard | September | RS | SS |
| HH_HH_01_07 | HH | graveyard | September | SS | SS |
| HH_HH_01_08 | HH | graveyard | September | RS | SS |
| HH_HH_01_09 | HH | graveyard | September | SS | SS |
| HH_HH_01_11 | HH | graveyard | September | RS | SS |
| HH_HH_01_13 | HH | graveyard | September | RS | SS |
| HH_HH_01_14 | HH | graveyard | September | RS | SS |
| HH_HH_01_16 | HH | graveyard | September | RS | SS |
| HH_HH_01_17 | HH | graveyard | September | SS | SS |
| SH_RZ_02_01 | RZ | graveyard | October | SS | SS |
| SH_RZ_02_02 | RZ | graveyard | October | SS | SS |
| SH_RZ_02_07 | RZ | graveyard | October | SS | SS |
| SH_RZ_02_08 | RZ | graveyard | October | SS | SS |
| SH_RZ_02_09 | RZ | graveyard | October | SS | SS |
| SH_RZ_02_10 | RZ | graveyard | October | SS | SS |
| SH_RZ_02_15 | RZ | graveyard | October | SS | SS |
| SH_RZ_02_16 | RZ | graveyard | October | SS | SS |
| SH_RZ_02_17 | RZ | graveyard | October | SS | SS |
| SH_RZ_02_20 | RZ | graveyard | October | SS | SS |
| HE_LM_02_05 | LM_1 | graveyard | September | SS | SS |
| HE_LM_02_07 | LM_1 | graveyard | September | SS | SS |
| HE_LM_02_09 | LM_1 | graveyard | September | SS | SS |
| HE_LM_02_14 | LM_1 | graveyard | September | SS | RS |
| HE_LM_02_16 | LM_1 | graveyard | September | SS | SS |
| HE_LM_03_03 | LM_1 | graveyard | September | RS | SS |
| HE_LM_03_08 | LM_1 | graveyard | September | RS | SS |
| HE_LM_03_10 | LM_1 | graveyard | September | RS | SS |
| HE_LM_03_14 | LM_1 | graveyard | September | SS | SS |
| HE_LM_03_21 | LM_1 | graveyard | September | RS | SS |
| NRW_VIE_01_04 | VIE_1 | graveyard | October | SS | SS |
| NRW_VIE_01_06 | VIE_1 | graveyard | October | SS | SS |
| NRW_VIE_02_05 | VIE_2 | graveyard | October | SS | SS |
| NRW_VIE_02_08 | VIE_2 | graveyard | October | SS | SS |
| NRW_VIE_02_10 | VIE_2 | graveyard | October | RS | SS |
| NRW_VIE_03_08 | VIE_2 | graveyard | October | RS | SS |
| NRW_VIE_03_11 | VIE_2 | graveyard | October | SS | SS |
| NRW_VIE_03_16 | VIE_2 | graveyard | October | RS | SS |
| NRW_VIE_04_04 | VIE_2 | graveyard | October | SS | SS |
| NRW_VIE_04_08 | VIE_2 | graveyard | October | RS | SS |
| BW_HD_01_09 | HD | graveyard | October | SS | SS |
| BW_HD_01_14 | HD | graveyard | October | failed | SS |
| BW_HD_01_17 | HD | graveyard | October | SS | SS |
| BW_HD_02_05 | HD | graveyard | October | SS | SS |
| BW_HD_02_11 | HD | graveyard | October | SS | SS |
| BW_HD_02_15 | HD | graveyard | October | SS | SS |
| BW_HD_02_17 | HD | graveyard | October | SS | SS |
| BW_HD_03_09 | HD | graveyard | October | RS | SS |
| BW_HD_03_14 | HD | graveyard | October | SS | SS |
| BW_HD_03_17 | HD | graveyard | October | SS | SS |
| WEI_1_01 | WEI | graveyard | September | SS | SS |
| WEI_1_02 | WEI | graveyard | September | SS | SS |
| WEI_1_03 | WEI | graveyard | September | SS | SS |
| WEI_1_04 | WEI | graveyard | September | SS | SS |
| WEI_1_05 | WEI | graveyard | September | SS | SS |
| WEI_1_06 | WEI | graveyard | September | SS | SS |
| WEI_1_07 | WEI | graveyard | September | SS | SS |
| WEI_1_08 | WEI | graveyard | September | SS | SS |
| WEI_1_09 | WEI | graveyard | September | SS | SS |
| WEI_1_10 | WEI | graveyard | September | SS | SS |
| WEI_1_11 | WEI | graveyard | September | SS | SS |
| WEI_1_12 | WEI | graveyard | September | SS | SS |
| WEI_1_13 | WEI | graveyard | September | SS | SS |
| WEI_1_14 | WEI | graveyard | September | SS | SS |
| WEI_1_15 | WEI | graveyard | September | SS | SS |
| WEI_1_16 | WEI | graveyard | September | SS | SS |
| WEI_1_17 | WEI | graveyard | September | SS | SS |
| WEI_1_18 | WEI | graveyard | September | SS | SS |
| WEI_1_19 | WEI | graveyard | September | SS | SS |
| SCH_2_03 | SCH | graveyard + paddock | September | failed | SS |
| SCH_2_10 | SCH | graveyard + paddock | September | SS | SS |
| SCH_2_12 | SCH | graveyard + paddock | September | SS | SS |
| SCH_2_16 | SCH | graveyard + paddock | September | SS | SS |
| SCH_2_17 | SCH | graveyard + paddock | September | failed | failed |
| SCH_2_18 | SCH | graveyard + paddock | September | SS | SS |
| WAL_3_09 | WAL | graveyard | August | SS | SS |
| GLA_2_07 | GLA | graveyard | September | failed | failed |
| GLA_2_10 | GLA | graveyard | September | failed | failed |
| GLA_2_12 | GLA | graveyard | September | SS | SS |
| GLA_2_13 | GLA | graveyard | September | SS | SS |
| GLA_2_14 | GLA | graveyard | September | SS | RS |
| GLA_2_15 | GLA | graveyard | September | SS | SS |
| GLA_2_16 | GLA | graveyard | September | SS | SS |
| GLA_3_20 | GLA | graveyard | September | SS | SS |
| HOF_2_10 | HOF | graveyard | September | SS | SS |
| HOF_2_12 | HOF | graveyard | September | SS | SS |
| HOF_2_17 | HOF | graveyard | September | SS | SS |
| HOF_2_19 | HOF | graveyard | September | SS | SS |
| HOF_3_01 | HOF | graveyard | September | RS | SS |
| HOF_3_06 | HOF | graveyard | September | SS | SS |
| HOF_3_11 | HOF | graveyard | September | SS | SS |
| HOF_3_12 | HOF | graveyard | September | RS | SS |
| HOF_3_13 | HOF | graveyard | September | RS | SS |
| HOF_3_14 | HOF | graveyard | September | SS | SS |
| HOF_3_17 | HOF | graveyard | September | SS | SS |
| HOF_3_19 | HOF | graveyard | September | SS | SS |
| WES_1_01 | WES | botanical garden | September | failed | failed |
| WES_1_02 | WES | botanical garden | September | SS | SS |
| WES_1_04 | WES | botanical garden | September | SS | SS |
| WES_1_11 | WES | botanical garden | September | SS | SS |
| WES_2_05 | WES | botanical garden | September | RS | failed |
| WES_2_10 | WES | botanical garden | September | SS | SS |
| WES_2_13 | WES | botanical garden | September | SS | failed |
| WES_2_19 | WES | botanical garden | September | SS | SS |
| WES_3_02 | WES | botanical garden | September | SS | SS |
| WES_3_06 | WES | botanical garden | September | SS | SS |
| WES_3_08 | WES | botanical garden | September | SS | SS |
| WES_3_13 | WES | botanical garden | September | SS | SS |
| WES_3_14 | WES | botanical garden | September | SS | SS |
| WES_3_18 | WES | botanical garden | September | RS | RS |
| SOS_1_05 | SOS | graveyard | August | RS | SS |
| SOS_1_16 | SOS | graveyard | August | SS | SS |
| SOS_1_17 | SOS | graveyard | August | SS | SS |
| SOS_1_18 | SOS | graveyard | August | SS | RS |
| SOS_3_01 | SOS | graveyard | August | SS | SS |
| SOS_3_07 | SOS | graveyard | August | SS | SS |
| SOS_3_09 | SOS | graveyard | August | SS | SS |
| SOS_3_16 | SOS | graveyard | August | SS | SS |
| SOS_3_17 | SOS | graveyard | August | SS | SS |
| NOR_1_01 | NOR | graveyard | September | RS | SS |
| NOR_1_05 | NOR | graveyard | September | RS | SS |
| NOR_1_06 | NOR | graveyard | September | SS | SS |
| NOR_1_07 | NOR | graveyard | September | RR | SS |
| NOR_1_14 | NOR | graveyard | September | RS | SS |
| NOR_1_18 | NOR | graveyard | September | RS | SS |
| HE_FB_33_04 | FB_3 | graveyard | July | SS | SS |
| BY_KT_70_02 | KT_1 | graveyard | August | SS | SS |
| BY_KT_70_20 | KT_1 | graveyard | August | SS | SS |
| BY_KT_71_17 | KT_2 | garden | August | SS | SS |
| BY_KT_71_18 | KT_2 | garden | August | SS | SS |
| BY_KT_72_04 | KT_2 | garden | August | SS | SS |
| BY_KT_72_14 | KT_2 | garden | August | SS | SS |
| BY_KT_90_06 | KT_3 | graveyard | August | SS | failed |
| BY_N_73_12 | N | graveyard | August | failed | failed |
| BY_N_73_13 | N | graveyard | August | failed | failed |
| BY_N_73_15 | N | graveyard | August | failed | SS |
| BY_N_73_17 | N | graveyard | August | SS | failed |
| BY_N_74_7 | N | graveyard | August | failed | SS |
| BY_N_74_14 | N | graveyard | August | SS | SS |
| BY_N_74_16 | N | graveyard | August | failed | SS |
| BY_N_74_19 | N | graveyard | August | failed | failed |
| BY_CO_86_09 | CO | graveyard | August | RS | SS |
| BY_CO_87_17 | CO | graveyard | August | failed | SS |
| BY_BA_92_06 | BA_1 | graveyard | August | RS | SS |
| BY_BA_94_06 | BA_1 | graveyard | August | failed | SS |
| BY_BA_94_10 | BA_1 | graveyard | August | failed | failed |
| BY_BA_94_11 | BA_1 | graveyard | August | failed | failed |
| SN_MW_104_06 | MW | garden | August | SS | SS |
| SN_MW_105_01 | MW | garden | August | SS | SS |
| SN_MW_105_04 | MW | garden | August | SS | SS |
| SN_MW_105_09 | MW | garden | August | SS | SS |
| SN_MW_106_20 | MW | garden | August | RS | failed |
| SN_L_119_03 | LB | garden | August | SS | SS |
| SN_L_119_08 | LB | garden | August | SS | SS |
| SN_L_119_10 | LB | garden | August | SS | SS |
| BY_FS_129_01 | FS | graveyard | September | failed | SS |
| BY_FS_129_02 | FS | graveyard | September | SS | failed |
| BY_ER_101_10 | ER | graveyard | August | SS | failed |
| BY_ER_101_12 | ER | graveyard | August | SS | SS |
| BW_RA_130_01 | RA | graveyard | October | SS | RS |
| BW_RA_131_05 | RA | graveyard | October | failed | failed |
| BW_RA_131_15 | RA | graveyard | October | SS | SS |
| BW_RA_131_20 | RA | graveyard | October | failed | failed |
| NRW_MS_135_10 | MS | graveyard | September | SS | SS |
| NRW_MS_135_11 | MS | graveyard | September | SS | SS |
| NRW_MS_135_18 | MS | graveyard | September | SS | SS |
| BW_HD_01_12 | HD | graveyard | October | failed | SS |
| SN_L_108_12 | LS | graveyard | August | SS | SS |
| SN_L_109_05 | LS | graveyard | August | SS | SS |
| SN_L_115_06 | LC | graveyard | August | RS | failed |
| SN_L_117_07 | LC | graveyard | August | RS | SS |

Table 3: Overview of the Cx. pipiens f. pipiens samples used for CO1 haplotype analysis.

| **Sample_ID** | **Sampling site abbreviation** | **Sampling site** | **Life stage sampled** |
| --- | --- | --- | --- |
| 7_11_A8 | RI | Rietschen | Egg |
| 8_11_F4 | BH | Breungeshain | Egg |
| 8_11_D9 | RB | Reinborn | Egg |
| 19_09_B9 | RB | Reinborn | Egg |
| 7_11_G3 | MF | Berlin-Marienfelde | Egg |
| NRW_VIE_01_04 | VIE | Kempen | Larvae |
| 7_11_A4 | RI | Rietschen | Egg |
| 7_11_D7 | BM | Berlin-Mitte | Egg |
| 8_11_A3 | FM | Frankfurt-Westend | Egg |
| SN_L_112_03 | L | Leipzig | Larvae |
| SH_RZ_02_07 | RZ | Mölln | Larvae |
| NRW_MS_135_03 | MS | Münster | Larvae |
| NI_FRI_46_15 | FRI | Varel | Larvae |
| NI_FRI_46_01 | FRI | Varel | Larvae |
| HE_FB_33_15 | FB | Bad Vilbel | Larvae |
| HE_FB_33_10 | FB | Bad Vilbel | Larvae |
| BY_WU_78_04 | WU | Würzburg | Larvae |
| HE_FB_09_16 | FB | Bad Vilbel | Larvae |
| BY_CO_86_03 | CO | Coburg | Larvae |
| BY_WU_78_16 | WU | Würzburg | Larvae |
| 7_11_H5 | MF | Berlin-Marienfelde | Egg |
| 2_9_F8 | BH | Breungeshain | Egg |
| 8_11_C12 | RB | Reinborn | Egg |
| 7_11_G5 | MF | Berlin-Marienfelde | Egg |
| 8_11_A2 | FM | Frankfurt-Westend | Egg |
| 8_11_A4 | FM | Frankfurt-Westend | Egg |
| BY_ER_101_14 | ER | Erlangen | Larvae |
| 19_09_B10 | RB | Reinborn | Egg |
| BW_HD_01_09 | HD | Heidelberg | Larvae |
| BW_RA_133_13 | RA | Muggensturm | Larvae |
| BY_ER_101_11 | ER | Erlangen | Larvae |
| BW_HD_01_17 | HD | Heidelberg | Larvae |
| BY_WU_78_03 | WU | Würzburg | Larvae |
| NRW_MS_135_14 | MS | Münster | Larvae |
| BW_HD_03_09 | HD | Heidelberg | Larvae |
| BY_ER_101_17 | ER | Erlangen | Larvae |
| BY_KT_70_16 | KT | Wiesentheid | Larvae |
| BY_WU_78_02 | WU | Würzburg | Larvae |
| SN_L_112_06 | L | Leipzig | Larvae |
| BW_KN_52_10 | KN | Gaienhofen | Larvae |
| BW_RA_133_11 | RA | Muggensturm | Larvae |
| BY_N_74_11 | N | Neurenberg | Larvae |
| BY_NM_76_01 | NM | Pyrbaum | Larvae |
| BY_NM_76_05 | NM | Pyrbaum | Larvae |
| BY_NM_76_06 | NM | Pyrbaum | Larvae |
| BY_NM_76_08 | NM | Pyrbaum | Larvae |
| BY_CO_86_17 | CO | Coburg | Larvae |
| NRW_VIE_02_05 | VIE | Kempen | Larvae |
| BY_N_75_15 | N | Neurenberg | Larvae |
| BY_WU_78_08 | WU | Würzburg | Larvae |
| BY_WU_78_12 | WU | Würzburg | Larvae |
| BY_WU_78_15 | WU | Würzburg | Larvae |
| HE_FB_33_07 | FB | Bad Vilbel | Larvae |
| HH_HH_01_14 | HH | Hamburg | Larvae |
| NRW_MS_134_04 | MS | Münster | Larvae |
| SN_L_113_07 | L | Leipzig | Larvae |
| BY_NM_76_13 | NM | Pyrbaum | Larvae |
| BY_WU_78_11 | WU | Würzburg | Larvae |
| BY_WU_78_13 | WU | Würzburg | Larvae |
| HE_FB_33_02 | FB | Bad Vilbel | Larvae |
| HE_LM_03_14 | LM | Bad Camberg | Larvae |
| HH_HH_01_13 | HH | Hamburg | Larvae |
| HH_HH_01_16 | HH | Hamburg | Larvae |
| NRW_MS_135_21 | MS | Münster | Larvae |
| SN_L_111_10 | L | Leipzig | Larvae |
| NRW_MS_135_08 | MS | Münster | Larvae |
| 7_11_E5 | BM | Berlin-Mitte | Egg |
| 8_11_B3 | FM | Frankfurt-Westend | Egg |
| SN_L_111_12 | L | Leipzig | Larvae |
| HH_HH_01_17 | HH | Hamburg | Larvae |
| HH_HH_01_11 | HH | Hamburg | Larvae |
| HH_HH_01_09 | HH | Hamburg | Larvae |
| HE_LM_03_10 | LM | Bad Camberg | Larvae |
| BY_NM_76_11 | NM | Pyrbaum | Larvae |
| BY_WU_78_01 | WU | Würzburg | Larvae |
| BY_N_75_17 | N | Neurenberg | Larvae |
| BY_ER_101_19 | ER | Erlangen | Larvae |
| BY_CO_86_10 | CO | Coburg | Larvae |
| 19_09_F5 | BM | Berlin-Mitte | Egg |
| 19_09_E10 | RI | Rietschen | Egg |
| 19_9_D1 | MF | Berlin-Marienfelde | Egg |
| 7_11_D3 | BM | Berlin-Mitte | Egg |
| 2_09_24_F9 | BH | Breungeshain | Egg |
| 7_11_A5 | RI | Rietschen | Egg |

Supplement Table 4: Mortality results of the CDC bottle bioassay for Cx. pipiens f. pipiens. PR = possible resistance.

| **Species** | **Insecticide (Permethrin: 43 μg/ml)** | | | **Control (acetone)** | |
| --- | --- | --- | --- | --- | --- |
|  | **No. of mosquitoes** | **Mortality (%)** | **Status** | **No. of Mosquitoes** | **Mortality (%)** |
| *Cx. pipiens* f*. pipiens* | 98 | 91.8% | PR | 29 | 0 |

Supplement Table 5: Targeted generalised additive model (GAM) screening results.

| **Response** | **Predictor** | **p_value** | **edf** | **Relationship** |
| --- | --- | --- | --- | --- |
| allele_frequency_perc | bio12 | 0.002347119 | 1.000136 | Linear |
| allele_frequency_perc | bio10 | 0.004616553 | 1.000159 | Linear |
| allele_frequency_perc | smod_class | 0.031411979 |  | Factor |

Supplement Table 6: The coefficients and smooth terms of the best-supported final GAMs.

| **Std_Error** | **edf** | **Test_Statistic** | **Statistic_Type** | **p_value** |
| --- | --- | --- | --- | --- |
| 0.136797 |  | 3.885 | t | 0.000533 |
| 0.004544 |  | 3.859 | t | 0.000572 |
| 0.140205 |  | -1.301 | t | 0.203208 |
| 0.144422 |  | -0.157 | t | 0.876301 |
| 0.167361 |  | -2.458 | t | 0.02004 |
| 0.105474 |  | -0.192 | t | 0.848703 |
| 0.149361 |  | 1.593 | t | 0.121779 |
| 0.288832 |  | 3.039 | t | 0.004929 |
|  | 3.43 | 8.43 | F | 0.000113 |
| 0.084162 |  | 10.69 | t | 6.36E-12 |
| 0.09586 |  | -1.373 | t | 0.18 |
| 0.092458 |  | -1.12 | t | 0.271 |
| 0.11306 |  | -1.105 | t | 0.278 |
| 0.072547 |  | 0.051 | t | 0.959 |
| 0.10267 |  | 0.761 | t | 0.453 |
| 0.194989 |  | -0.809 | t | 0.425 |
|  | 2.997 | 2.476 | F | 0.0601 |
|  | 1 | 12.65 | F | 0.00131 |
| 6.8866 |  | 0.972 | t | 0.339 |
| 6.8372 |  | 0.017 | t | 0.987 |
| 7.2004 |  | 1.578 | t | 0.125 |
| 8.172 |  | 0.846 | t | 0.404 |
| 4.7453 |  | 0.039 | t | 0.969 |
| 7.0178 |  | -0.389 | t | 0.7 |
| 12.7169 |  | 0.025 | t | 0.98 |

Supplement Table 7: Comparison of the candidate GAMs using AIC and AICc.

| **AIC** | **AICc** | **delta** | **weight** | **Best_Supported** |
| --- | --- | --- | --- | --- |
| 18.13654 | 30.9 | 0 | 0.992 | Yes |
| 20.85481 | 41.1 | 10.16 | 0.006 | No |
| 33.06906 | 43.5 | 12.55 | 0.002 | No |
| 34.21298 | 51 | 20.01 | 0 | No |
| -15.21204 | -5.1 | 0 | 0.972 | Yes |
| -12.63557 | 2 | 7.07 | 0.028 | No |
| **AIC** | **AICc** | **delta** | **weight** | **Best_Supported** |
| 296.415 | 303.1 | 0 | 1 | Yes |
| 302.331 | 325.2 | 22.11 | 0 | No |
| 304.893 | 337.1 | 34.02 | 0 | No |
| 303.423 | 341.4 | 38.34 | 0 | No |

Supplement Table 8: Diagnostic statistics for the best-supported GAMs.

| **Best-supported model** | **k-index** | **k-check p** | **Shapiro-Wilk W** | **Shapiro p** | **Breusch-Pagan BP** | **BP p** | **Concurvity smooth (estimate)** | **Concurvity smooth (observed)** | **Concurvity smooth (worst)** |
| --- | --- | --- | --- | --- | --- | --- | --- | --- | --- |
| model_kdr_precip_controls_nospace | 0.96 | 0.44 | 0.96818 | 0.3612 | 10.603 | 0.001129 | 0.125 | 0.136 | 0.346 |
